# Endothelial cell response to Hedgehog ligands depends on their processing

**DOI:** 10.1101/2020.03.03.974444

**Authors:** Pierre-Louis Hollier, Candice Chapouly, Aissata Diop, Sarah Guimbal, Lauriane Cornuault, Alain-Pierre Gadeau, Marie-Ange Renault

**Author notes:** **Corresponding author** Marie-Ange Renault Inserm U1034, 1, avenue de Magellan, 33604 Pessac, France. These authors contributed equally to this work (Pierre-Louis Hollier, Candice Chapouly).

## Abstract

**Rationale:** The therapeutic potential of Hedgehog (Hh) signaling agonists for vascular diseases is of growing interest. However, molecular and cellular mechanisms underlying the role of the Hh signaling in vascular biology remain poorly understood.

**Objective:** The purpose of the present paper is to clarify some conflicting literature data.

**Findings:** With this goal we have demonstrated that, unexpectedly, ectopically administered N-terminal Sonic Hedgehog (N-Shh) and endogenous endothelial-derived Desert Hedgehog (Dhh) induce opposite effects in endothelial cells. (ECs). Notably, endothelial Dhh acts under its full-length soluble form (FL-Dhh) and activates Smoothened in ECs, while N-Shh inhibits it. At molecular level, N-Shh prevents FL-Dhh binding to Patched-1 demonstrating that N-Shh acts as competitive antagonist to FL-Dhh. Besides, we found that even though FL-Hh ligands and N-Hh ligands all bind Patched-1, they induce specific Patched-1 localization. Finally, we confirmed that in a pathophysiological setting i.e. brain inflammation, astrocyte-derived N-Shh act as a FL-Dhh antagonist.

**Conclusion:** The present study highlights for the first time that FL-Dhh and N-Hh ligands have antagonistic properties especially in ECs, and demonstrates that Hh ligands or forms of Hh ligands cannot be used instead of another for therapeutic purposes.

---

Endothelium integrity is essential to vascular homeostasis, since a failure of this system represent a critical factor in cardiovascular and cerebrovascular disease pathogenesis. Indeed, the endothelium is involved in many physiological processes such as angiogenesis, vascular permeability, vascular tone regulation, blood coagulation as well as homing of immune cells to specific sites of the body. Conversely, endothelial dysfunction is associated with excessive vasoconstriction especially because of impaired endothelial nitric oxide (NO) production. Also, it is characterized by abnormal vascular leakage due to altered endothelial intercellular junctions. Finally, dysfunctional endothelial cells (ECs) acquire pro-inflammatory and pro-thrombotic phenotypes by expressing increased levels of adhesion and pro-thrombotic molecules such as vascular cell adhesion molecule-1 (VCAM-1) and intercellular adhesion molecule-1 (ICAM-1).

Evidences accumulated within the past decades, identified Hedgehog (Hh) signaling as a new regulator of vascular homeostasis. More specifically, It has been shown to regulate both angiogenesis ^1^ and micro-vessel integrity ^2^. Indeed, previous investigations have reported that inhibition of Hh protein activity, by neutralizing 5E1 antibody administration, impairs ischemia-induced angiogenesis both in the setting of hind limb ischemia and myocardial infarction in mice ^3,4^. Accordingly, when Sonic Hedgehog (Shh), one of the Hh ligands, was administered either as a recombinant protein or via gene therapy, it promoted neovascularization of ischemic tissues by promoting both angiogenesis ^1^ and endothelial progenitor cell recruitment ^5^. Besides, Hh signaling was shown to promote blood brain barrier integrity and immune quiescence both in the setting of multiple sclerosis ^2^ and in the setting of stroke ^6^. Additionally we have shown that disruption of Hh signaling specifically in ECs induces blood-nerve barrier breakdown and peripheral nerve inflammation ^7^. As a consequence, the therapeutic potential of Hh signaling agonists for vascular diseases is of growing interest ^8^ ^9^ ^10^ ^11^ ^12^ ^13^. However, molecular and cellular mechanisms underlying the role of the Hh ligands in vascular biology remain poorly understood and conflicting results have been reported ^14^. For Instance, while, as described above, most studies agree in reporting that Shh is proangiogenic ^1,5,15^, unexpectedly, ischemia-induced angiogenesis was shown to be accelerated in Shh deficient mice ^16^. Similarly, while Hh signaling is typically believed to promote endothelial barrier integrity ^2,17^, Glioblastoma-derived Desert Hedgehog (Dhh) was shown to disrupt the BBB ^18^. Additionally, an early study reported that ectopic Shh overexpression in the dorsal neural tube induces hemorrhage in the spinal cord ^19^.

The Hedgehog (Hh) family of morphogens, which was identified nearly 4 decades ago in drosophila as crucial regulators of cell fate determination during embryogenesis, includes three members: Shh, Indian hedgehog (Ihh) and Dhh ^20^. The interaction of Hh proteins with their specific receptor Patched-1 (Ptch1) de-represses the transmembrane protein Smoothened (Smo), which activates downstream pathways, including the Hh canonical pathway leading to the activation of Gli family zinc finger (Gli) transcription factors and so-called Hh non canonical pathways, which are independent of Smo and/or Gli ^21^. However, while Shh, Ihh and Dhh, all bind to the same receptor, Ptch1, with close affinities, their ability to activate Gli-dependent transcription, in fibroblast cell lines is dramatically different. Ihh is 6 times less potent than Shh, and Dhh is more than 10 times less potent than Ihh ^22^. This has been one of the reasons why Shh is the main Hh gene considered for therapy. In contrast, some of Hh biological effects including Islet-1 induction ^22^ or activation of RhoA in ECs are equally induced by the 3 proteins ^23^.

Besides, Shh is synthetized as a pre-protein whom signal sequence is first cleaved to produce a full length unmodified form. Then an autocatalyic reaction removes the carboxy-terminal domain and attaches a cholesterol moiety to the newly exposed carboxy-terminus. Shh is further modified by Hedgehog acyltransferase (Hhat) which catalyzed the addition of a palmitate to the amino-terminus ^24^ to generate the Shh active form (N-Shh). Secretion and solubility of cholesterol-modified Shh depend on the transmembrane protein Disp1 (dispatched RND transporter family member 1) and the cell surface protein Scube2 (signal peptide, CUB domain and EGF like domain containing 2) ^25^. Both Disp1 and Scube2 bind the cholesterol-anchor of Shh. On the contrary to Shh, Ihh and Dhh processing have been poorly investigated, and may differ. Indeed, Dhh is suggested not to undergo efficient autocatalytic cleavage ^26^,

The purpose of the present paper is to compare the vascular effects of endothelial-derived Dhh *versus* ectopically administered N-Shh in order to determine whether Hh ligands rather promote vessel integrity and quiescence or destabilize vessels to promote angiogenesis and to validate some debated vascular effects of Hh ligands.

## Results

### Ectopic administration of N-Shh promotes angiogenesis while endothelial Dhh inhibits it

Since Hh ligands are well known for their pro-angiogenic effect ^1^, we first used the mouse corneal angiogenesis model to compare the effect of ectopically administered recombinant N-Shh (rec N-Shh) and endogenous endothelial Dhh on angiogenesis. Consistent with previous investigations ^1,15^, implantation of N-Shh-containing pellets in the cornea of mice induced angiogenesis (Figure 1A-B). However, when VEGFA-containing pellets were implanted in the corneas of both EC specific Dhh knockout mice (Dhh^ECKO^) and their control littermate, angiogenesis was significantly enhanced in Dhh^ECKO^ mice compared to control mice (Figure 1C-D). This later result indicates that unlike ectopically administered rec N-Shh, endogenous endothelial Dhh is rather anti-angiogenic. Similar results were obtained when we quantified migration of cultured ECs. Indeed, while rec N-Shh protein promoted EC migration (Figure 1E), Dhh knock down in ECs also increased cell migration (Figure 1F). Altogether, this first set of data demonstrates for the first time that ectopic N-Shh and EC-derived endogenous Dhh induce opposite effects especially on ECs.

**Figure 1:**
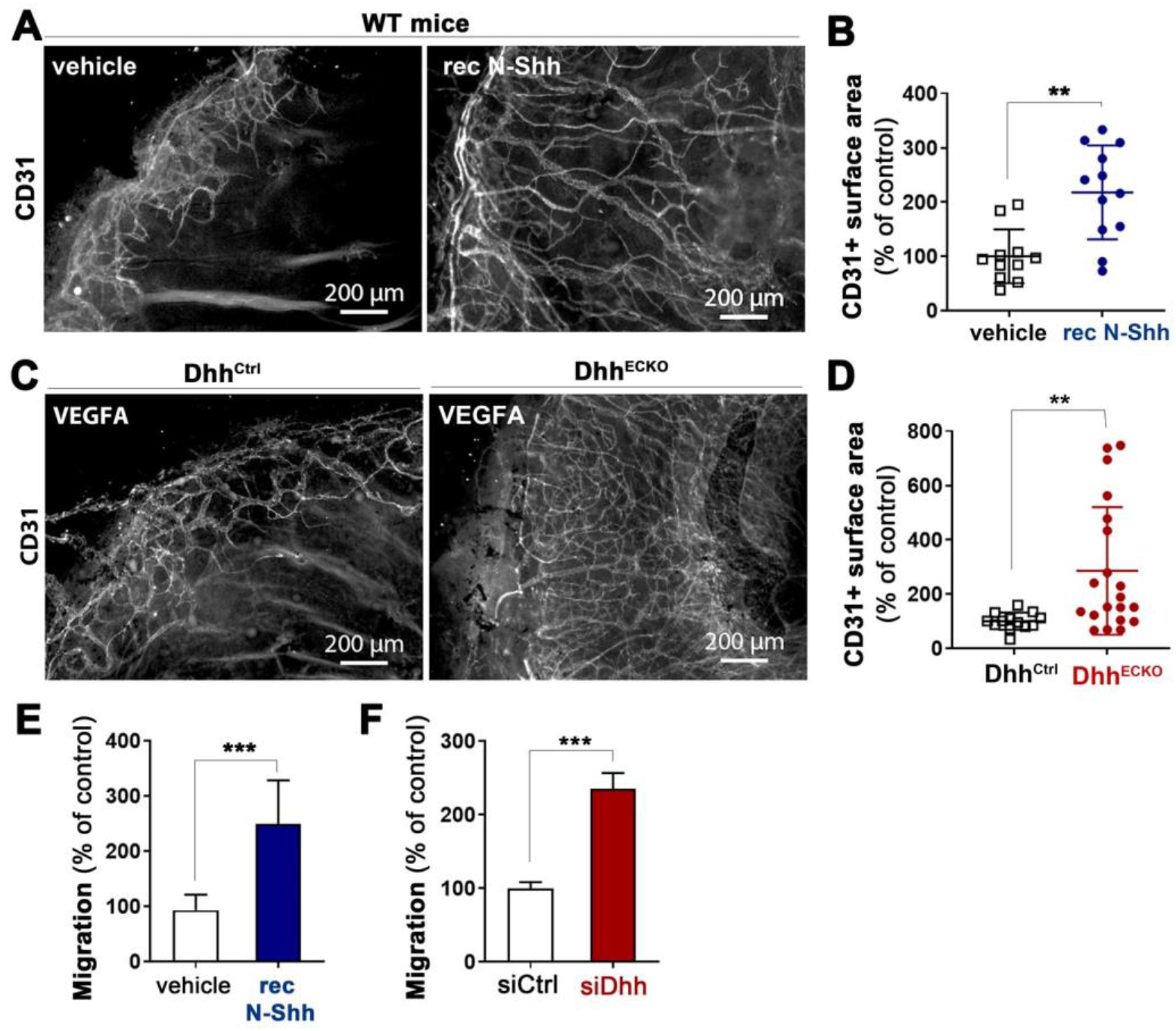
N-Shh promotes angiogenesis while EC-derived Dhh inhibits it. (**A-B**) recombinant N-Shh-containing or control pellets were implanted in the corneas of WT mice (n=12 and 11 corneas respectively). Mice were sacrificed 9 days later. (**A**) Whole mount corneas were immunostained with anti-CD31 antibodies to identify blood vessels. Representative pictures are shown. (**B**) Angiogenesis was quantified as the percentage of CD31+ surface area. (**C-D**) VEGFA containing pellets were implanted in the corneas of Cdh5-Cre^ERT2^ Dhh^Flox/Flox^ (Dhh^ECKO^) and Dhh^Flox/Flox^ (Dhh^Ctrl^) mice 2 week after they were administered with tamoxifen (n=20 and 13 corneas respectively). (**C**) Whole mount corneas were immunostained with anti-CD31 antibodies to identify blood vessels. Representative pictures are shown. (**D**) Angiogenesis was quantified as the percentage of CD31+ surface area. (**E**) HUVEC migration was assessed in a chemotaxis chamber in the presence or not of 1 μg/mL recombinant N-Shh. The experiment was repeated 3 times, each experiment included n=4 wells/conditions. (**F**) HUVECs were transfected with Dhh or control siRNAs. Cell migration was assessed in a chemotaxis chamber. The experiment was repeated 3 times, each experiment included n=4 wells/conditions. **: p≤0.01; ***: p≤0.001. Mann Whitney test

### Endothelial Dhh promotes BBB integrity while Ectopic administration of N-Shh disrupts it

To confirm that endothelial Dhh and ectopic N-Shh induce opposite effects on ECs, we compared their effect on endothelial barrier integrity. Consistent, with previous investigations ^17,27^, expression of both Cdh5 and Cldn5 was significantly diminished at the BBB of Dhh^ECKO^ mice compared to littermate controls (Figure 2A-D). This was associated with fibrinogen and albumin extravasation (Figure 2A-B, E-F) demonstrating abnormal BBB permeability. To investigate the effects of N-Shh, both N-Shh-encoding and control lentivirus were administered locally in the cortex of mice. As shown in Figure 2G-H, N-Shh significantly reduced Cdh5 expression at the BBB which was associated with increased albumin and fibrinogen extravasation (Figure 3G, I-J). Consistent results were obtained in HUVECs in which both Dhh knock down or treatment with recombinant N-Shh protein increased Cdh5 junction thickness indicating destabilization of adherens junctions (Supplementary Figure 1A-D). Additionally, these results were confirmed in mouse primary cultured brain ECs that were either prepared from Dhh^ECKO^ mice and littermate controls, or from WT mice which were then treated with recombinant N-Shh protein versus BSA. As shown in Supplementary Figure 1E-F Dhh deficiency in brain ECs led to disorganized Cdh5 junctions and decreased expression of tight junction proteins including Cldn5 and Ocln. Treatment with rec N-Shh also altered brain EC junctions as it led to increased Cdh5 and Cldn5 internalization (Supplementary Figure 1G-H).

**Figure 2:**
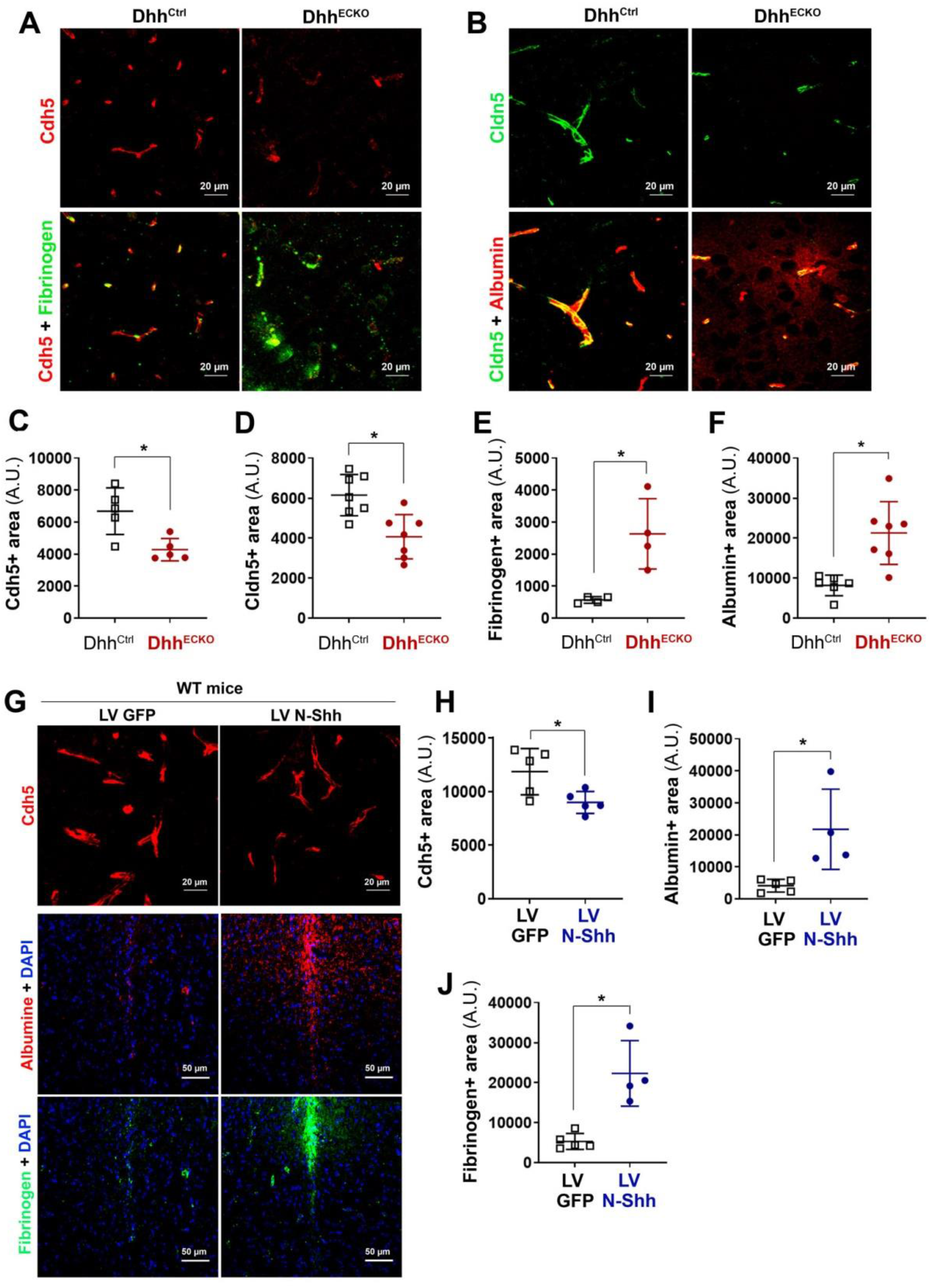
EC-derived Dhh prevents BBB opening while N-Shh disrupts it. (**A-F**) Cdh5-Cre^ERT2^ Dhh^Flox/Flox^ (Dhh^ECKO^) and Dhh^Flox/Flox^ (Dhh^Ctrl^) mice were sacrificed 2 weeks after they were administered with tamoxifen (n=7 and 6 mice respectively). (**A-B**) Brain sagittal sections were immunostained with either stained with anti-Cdh5 (in red), and anti-Fibrinogen (in green) antibodies (**A**) or stained with anti-Cldn5 (in green) and anti-Albumin (in red) antibodies (**B**). Representative confocal images are shown. (**C**) Cdh5 expression was quantified as the Cdh5+ surface area. (**D**) Cldn5 expression was quantified as the Cldn5+ surface area. (**E**) Fibrinogen extravasation was quantified as the fibrinogen+ surface area. (**F**) Albumin extravasation was quantified as the albumin+ surface area. (**G-J**) WT mice were administered in the cerebral cortex with lentiviruses encoding N-Shh (LV N-Shh) or not (LV GFP) (n= 5 mice in each group). Mice were sacrificed 14 days later. (**G**) Brain sagittal sections, chosen at the injection site, were immunostained with either anti-Cdh5 (in red), anti-Albumin (in red) or anti-Fibrinogen (in green) antibodies. Representative confocal images are shown. (**H**) Cdh5 expression was quantified as the Cdh5+ surface area. (**I**) Albumin extravasation was quantified as the albumin+ surface area. (**J**) Fibrinogen extravasation was quantified as the fibrinogen+ surface area. *: p≤0.05; **: p≤0.01. Mann Whitney test.

**Figure 3:**
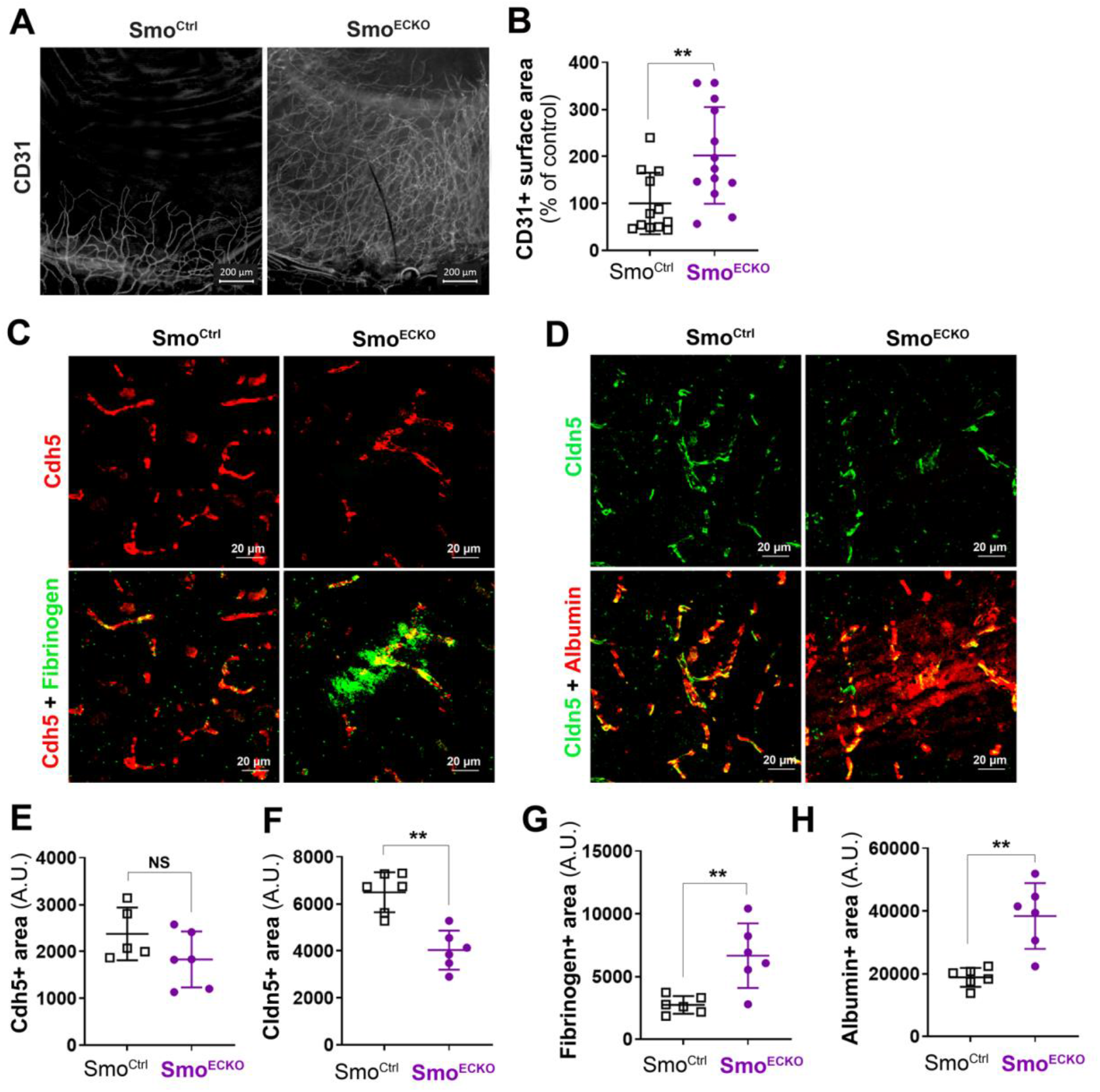
Inhibition of Hh signaling in ECs promotes angiogenesis and destabilizes the BBB. (**A-B**) VEGFA containing pellets were implanted in the corneas of Cdh5-Cre^ERT2^ Smo^Flox/Flox^ (Smo^ECKO^) and Smo^Flox/Flox^ (Smo^Ctrl^) mice 2 weeks after they were administered with tamoxifen (n=13 and 12 corneas respectively). (**A**) Whole mount corneas were immunostained with anti-CD31 antibodies to identify blood vessels. Representative pictures are shown. (**B**) Angiogenesis was quantified as the percentage of CD31+ surface area. (**C-H**) Smo^ECKO^ and Smo^Ctrl^ mice were sacrificed 2 weeks after they were administered with tamoxifen (n= 6 mice in each group). (**C-D**) Brain sagittal sections were immunostained with either anti-Cdh5 (in red), and anti-Fibrinogen (in green) antibodies (**C**) or anti-Cldn5 (in green) and anti-Albumin (in red) antibodies (**D**). Representative confocal images are shown. (**E**) Cdh5 expression was quantified as the Cdh5+ surface area. (**F**) Cldn5 expression was quantified as the Cldn5+ surface area. (**G**) Fibrinogen extravasation was quantified as the fibrinogen+ surface area. (**H**) Albumin extravasation was quantified as the albumin+ surface area. **: p≤0.01. Mann Whitney test.

This second set of data confirms that endothelial Dhh and ectopic N-Shh trigger opposite effects: while endothelial Dhh promotes endothelium integrity and prevents angiogenesis, ectopic N-Shh destabilizes blood vessels and promotes angiogenesis.

### Activating Smo mimics endothelial Dhh induced effects while inhibiting Smo recapitulates N-Shh effects

Since endothelial Dhh and ectopic N-Shh induce opposite effects in ECs, we hypothesized that one of the ligand would activate Hh signaling in EC while the other would inhibit it. To test this hypothesis, we assessed the functional consequences of Smo modulation since the classical read out to assess Hh signaling activity (i.e. Gli activity) is not modulated in ECs (Supplementary Figure 2). First, we investigated the role of Smo in angiogenesis; VEGFA containing pellets were implanted in the corneas of both Smo^ECKO^ mice and their littermate controls. Interestingly, Smo deficiency significantly increased VEGFA-induced angiogenesis (Figure 3A-B). Consistently, *in vitro*, Smo inhibition with GDC-0449 significantly stimulated HUVEC migration while Smo activation with the Smo agonist SAG didn’t (Supplementary Figure 3A). Then we investigated the role of Smo on endothelial barrier integrity; Smo deficiency in ECs significantly decreased both Cdh5 and Cldn5 expression at the BBB (Figure 3 C-F) which was associated with increased fibrinogen and albumin extravasation (Figure 3C-D, G-H). *In vitro*, GDC-0449 increased Cdh5 junction thickness (Supplementary Figure 3B-C).

Altogether these results suggest that Dhh activates Hh signaling in ECs via Smo while ectopic N-Shh inhibits it. In order to verify this hypothesis, we performed rescue experiments. ECs were either transfected with control or Dhh siRNA and then treated or not with SAG. As expected, the increased Cdh5 junction thickness observed in the absence of Dhh was reversed in the presence of SAG (Supplementary Figure 3D-E). In parallel, ECs were treated with rec N-Shh in the presence or not of SAG. Once again, N-Shh-induced increased Cdh5 junction thickness was reversed in the presence of SAG confirming that ectopic N-Shh inhibits Dhh-induced autocrine signaling in ECs.

### Dhh promotes endothelium integrity under its 43 kDa full-length soluble form

With the aim to investigate why endothelial Dhh and ectopic N-Shh induce opposite effects in ECs, we first wished to characterize under which form endothelial Dhh promotes endothelial integrity. Notably, while Shh is mainly processed as a 19 kDa N-terminal peptide and secreted as both a 45 kDa full length peptide and 19 kDa N-terminal peptide (Supplementary Figure 4A), Dhh is mainly produced as a 43 kDa full length protein and secreted exclusively under this form (Supplementary Figure 4B).

We then tested whether Dhh requires being palmitoylated using RU-SKI. RU-SKI did not modulate either HUVEC migration or HUVEC junction thickness (Figure 4A-C) indicating that Dhh does not need to be palmitoylated to promote endothelium integrity. To investigate the role of Disp1, a transmembrane protein necessary for Hh ligand transport across the plasma membrane, HUVECs were transfected with Disp1 or control siRNA. Disp1 knock down in HUVECs enhanced HUVEC migration (Figure 4D), increased Cdh5+ junction thickness (Figure 4E-F) and endothelium permeability (Figure 4G). Consistent with Dhh being exposed at the cell surface or secreted, Hh blocking antibodies increased Cdh5 junction thickness in HUVECs (Figure 4H-I). Finally, to investigate whether Dhh acts as a membrane bound protein or as a soluble secreted protein, we tested whether conditioned medium containing soluble 43 kDa Dhh would restore adherens junction integrity in Dhh siRNA transfected HUVECs. Soluble 43 kDa Dhh prevented both Dhh siRNA-induced Cdh5+ junction thickening (Figure 4J-K) and endothelial permeability (Figure 4L) demonstrating that Dhh promotes endothelial integrity as a soluble 43 kDa, full lengh.

**Figure 4:**
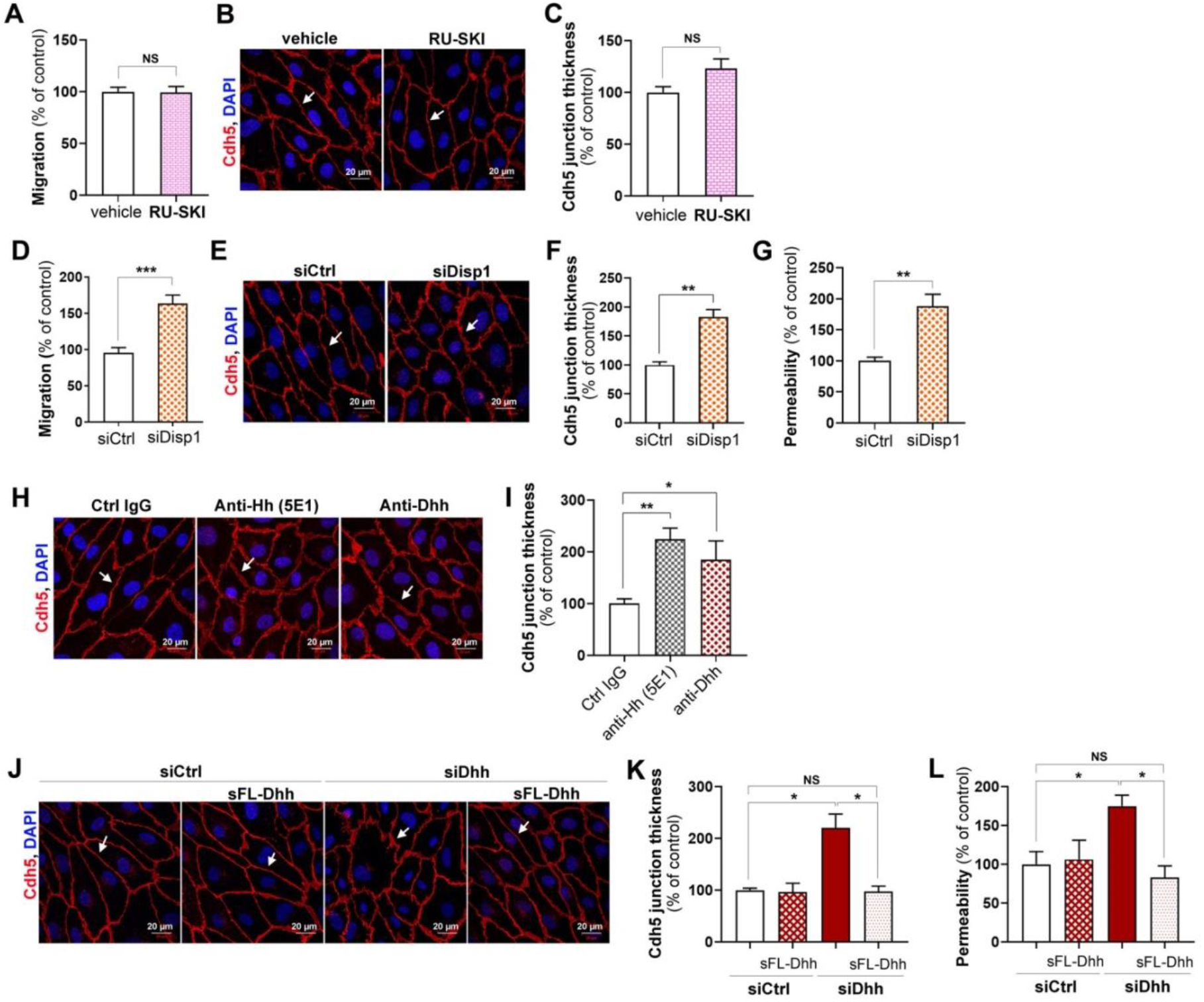
Dhh promotes endothelium integrity under its soluble form. (**A-C**) HUVECs were treated or not with 10 μmol/L RU-SKI (Sigma-Aldrich). (**A**) Cell migration was assessed in a chemotaxis chamber. The experiment was repeated 3 times, each experiment included n=4 wells/conditions. (**B**) Cdh5 localization was evaluated by immunofluorescent staining (in red) of a confluent cell monolayer and (**C**) quantified as the mean junction thickness using Image J software. The experiment was repeated at least 4 times. (**D-G**) HUVECs were transfected with Disp1 or control siRNAs. (**D**) Cell migration was assessed in a chemotaxis chamber. The experiment was repeated 3 times, each experiment included n=4 wells/conditions. (**E**) Cdh5 localization was evaluated by immunofluorescent staining (in red) of a confluent cell monolayer and (**F**) quantified as the mean junction thickness using Image J software. The experiment was repeated at least 4 times. (**G**) Endothelial monolayer permeability to 70 kDa FITC-dextran was assessed using Transwells. The experiment was repeated 3 times, each experiment included triplicates. (**H-I**) HUVECs were treated either with 2 μg/mL Hh blocking antibodies (5E1, DSHB), 2 μg/mL Dhh blocking antibodies (Santa-Cruz Biotechnology, sc271168), or 2 μg non-blocking mouse IgGs. (**H**) Cdh5 localization was evaluated by immunofluorescent staining (in red) of a confluent cell monolayer and (**I**) quantified as the mean junction thickness using Image J software. The experiment was repeated at least 4 times. (**J-L**) HUVECs were transfected with Dhh or control siRNAs, then treated with HeLa conditioned medium containing or not soluble FL-Dhh (sFL-Dhh). (**J**) Cdh5 localization was evaluated by immunofluorescent staining (in red) of a confluent cell monolayer and (**K**) quantified as the mean junction thickness using Image J software. The experiment was repeated at least 4 times. (**L**) Endothelial monolayer permeability to 70 kDa FITC-dextran was assessed using Transwells. The experiment was repeated 3 times, each experiment included triplicates. *: p≤0.05; **: p≤0.01; NS: not significant. Mann Whitney test or Kruskal-Wallis test followed by Dunn’s multiple comparison test.

In the meantime, we compared the effects of conditioned medium of HeLa cells transfected with either FL-Dhh, N-Shh, FL-Shh or N-Dhh encoding plasmids on HUVEC barrier properties. The conditioned medium from HeLa cells transfected with both N-Shh and FL-Shh encoding plasmids increased Cdh5 junction thickness (Supplementary Figure 4C-D) and increased endothelial monolayer permeability (Supplementary Figure 4E) confirming recombinant N-Shh protein effects. Notably, 17 kDa N-Dhh containing conditioned medium recapitulated the effects of N-Shh but not these of FL-Dhh demonstrating that Dhh requires being under its 43 kDa full length form to promote endothelium integrity and suggesting that any N-Hh ligands would inhibit Hh signaling in ECs.

### N-Shh prevents FL-Dhh binding to Ptch1

We hypothesized that N-Hh ligands inhibit Hh signaling in ECs by preventing Dhh-induced signaling. First, we quantified Dhh binding to Ptch1 in the presence or absence of N-Shh. To do so, we verified that both 43 kDa Dhh and N-Shh bind Ptch1 in immunoprecipitation assays (Figure 5A-B). We then measured the amount of 43 kDa Dhh bound to Ptch1 in the presence or absence of N-Shh by immunoprecipitation assay. Dhh binding to Ptch1 was reduced by about 70-80% in the presence of N-Shh (Figure 5C). The same result was obtained in the proximity ligation assay (Figure 5D-E). Notably, N-Dhh prevented 43 kDa Dhh binding to Ptch1 to the same extent as N-Shh while FL-Shh only decreased 43 kDa Dhh binding to Ptch1 by 50% (Supplementary Figure 5B).

**Figure 5:**
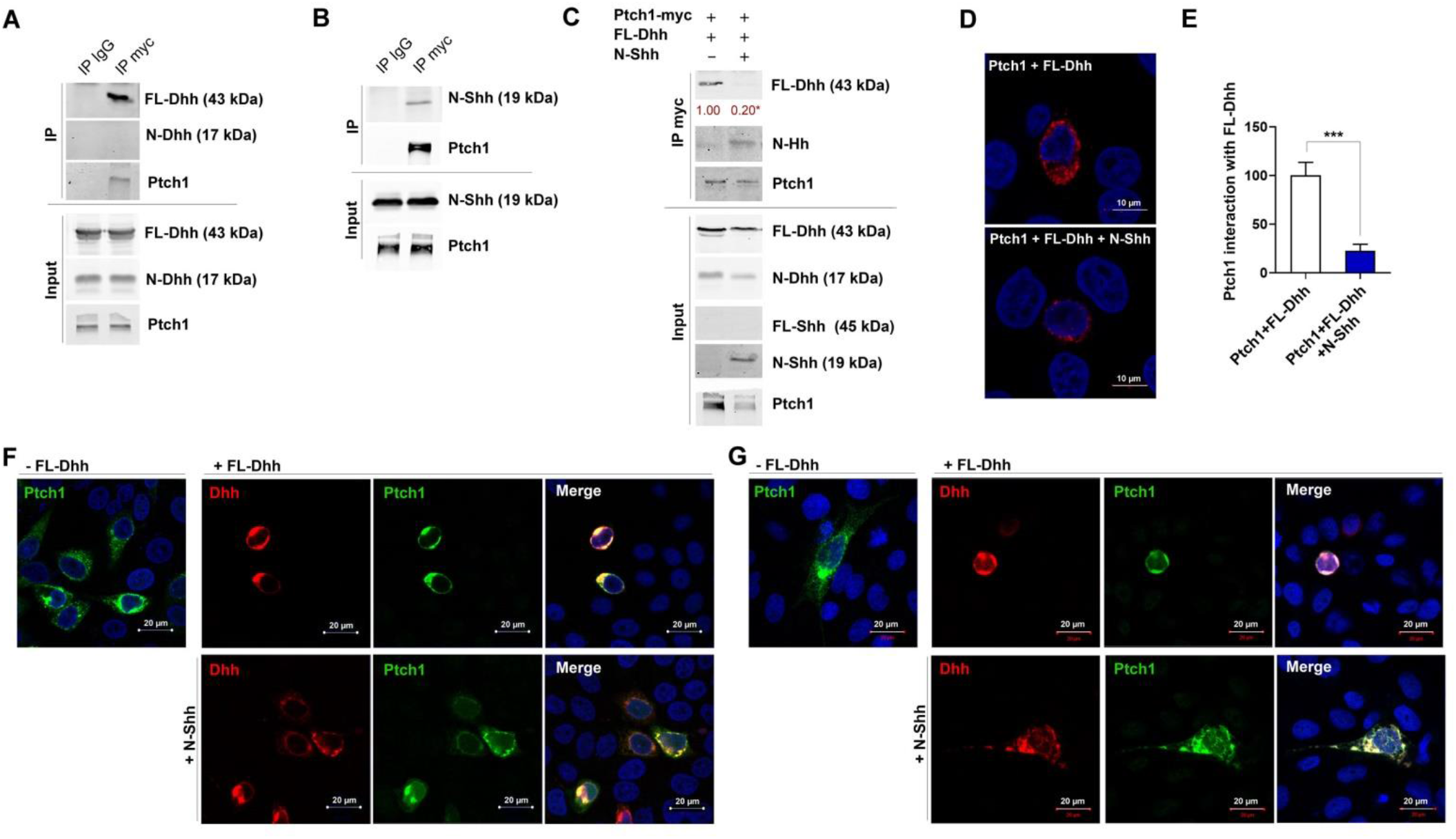
N-Shh prevents FL-Dhh binding to Ptch1. (**A**) HeLa cells were co-transfected with Ptch1-myc and FL-Dhh encoding plasmids. Dhh interaction with Ptch1 was assessed by co-immunoprecipitation assay. (**B**) HeLa cells were co-transfected Ptch1-myc and N-Dhh encoding plasmids. Dhh interaction with Ptch1 was assessed by co-immunoprecipitation assay. (**C-F**) HeLa cells were co-transfected Ptch1-myc and FL-Dhh encoding plasmids together with or without N-Shh encoding plasmids. (**C**) Dhh interaction with Ptch1 was assessed by co-immunoprecipitation assay. The experiment was repeated at least 4 times. (**D**) Dhh interaction with Ptch1 was assessed by proximity ligation assay and (**E**) quantified as mean red staining intensity/cell. The experiment was repeated at least 4 times. (**F**) Ptch1 and Dhh localization was assessed by immunostaining. (**G**) EA.hy926 cells were co-transfected with Ptch1-myc and FL-Dhh encoding plasmids together with or without N-Shh encoding plasmids. Ptch1 and Dhh localization was assessed by immunostaining. *: p≤0.05; **: p≤0.01; ***: p≤0.001; Mann Whitney test.

Altogether these results indicate for the first time that N-Hh ligands may act as competitive antagonists to 43 kDa Dhh in ECs.

### Each forms of Hh ligands induces a specific localization in Ptch1

The key remaining question was to determine why N-Hh ligands aren’t recapitulating 43 kDa Dhh-induced effects once they bind Ptch1. Therefore, we decided to investigate the earliest signaling events following Hh ligands binding to Ptch1. Upon Hh ligands binding to Ptch1, Ptch1 is being internalized and degraded by the ubiquitin proteasome system. We first verified that 43 kDa Dhh induces Ptch1 degradation through the proteasome. As shown in Supplementary Figure 6A, 43 kDa Dhh induces Ptch1 degradation just like the E3 ubiquitin ligase Itch ^28^ and Dhh-induced Ptch1 degradation is prevented in the presence of the proteasome inhibitor MG132. However, N-Shh, N-Dhh and FL-Shh all induced Ptch1 degradation (Supplementary Figure 6B) and neither of them modulated 43 kDa Dhh-induced Ptch1 degradation (Supplementary Figure 6C).

On the contrary, each Hh ligand and form of Hh ligand induced a specific localization of Ptch1. 43 kDa Dhh provoked a peri-nuclear localization of Ptch1, FL-Shh triggered Ptch1 endocytosis into vesicle distributed within the entire cell, while N-Shh and N-Dhh induced the formation of big Ptch1 aggregates in cells (Supplementary Figure 6D). As a consequence, N-Shh prevented 43 kDa Dhh-induced massive peri-nuclear localization of Ptch1 (Figure 5F). Finally, we verified whether the same was true in ECs, to this end, we repeated this last experiment in EA.hy926 cells. Consistently with results obtained in HeLa, N-Shh prevented 43 kDa Dhh-induced Ptch1 peri-nuclear localization (Figure 5G).

### Astrocyte-derived Shh disrupts BBB in the setting of brain inflammation

Finally, since we have only been investigating the vascular effects of ectopic N-Shh so far, we wished to investigate whether such regulation of Hh signaling exist in pathophysiological settings. Notably, Shh is barely expressed in the healthy vasculature; however, its expression is strongly upregulated in perivascular astrocytes upon brain inflammation (Supplementary Figure 7B) ^2^. More specifically, Shh mRNA is strongly up regulated in astrocytes upon Il-1β treatment (Supplementary Figure 7C). We performed a series of experiments to investigate what would be the paracrine effects of astrocyte-derived Shh on endothelial barrier integrity. Therefore, we first verified that Il1β-treated astrocytes produce a N-terminal fully processed Shh form (Supplementary Figure 7D). We then compared BBB integrity in the setting of Il1β-induced acute inflammation in both mice deficient for Shh in astrocytes (Shh^ACKO^) and in their control littermates. At 7 days after AdIl1β brain microinjection, expression of both Cdh5 and Cldn5 were significantly increased in the absence of astrocyte-derived Shh (Figure 6A-C). This was associated with decreased extravasation of serum proteins including fibrinogen and albumin (Figure 6A, D-E) demonstrating for the first time that astrocytic Shh expression disrupt BBB integrity during acute neuro-inflammation.

**Figure 6:**
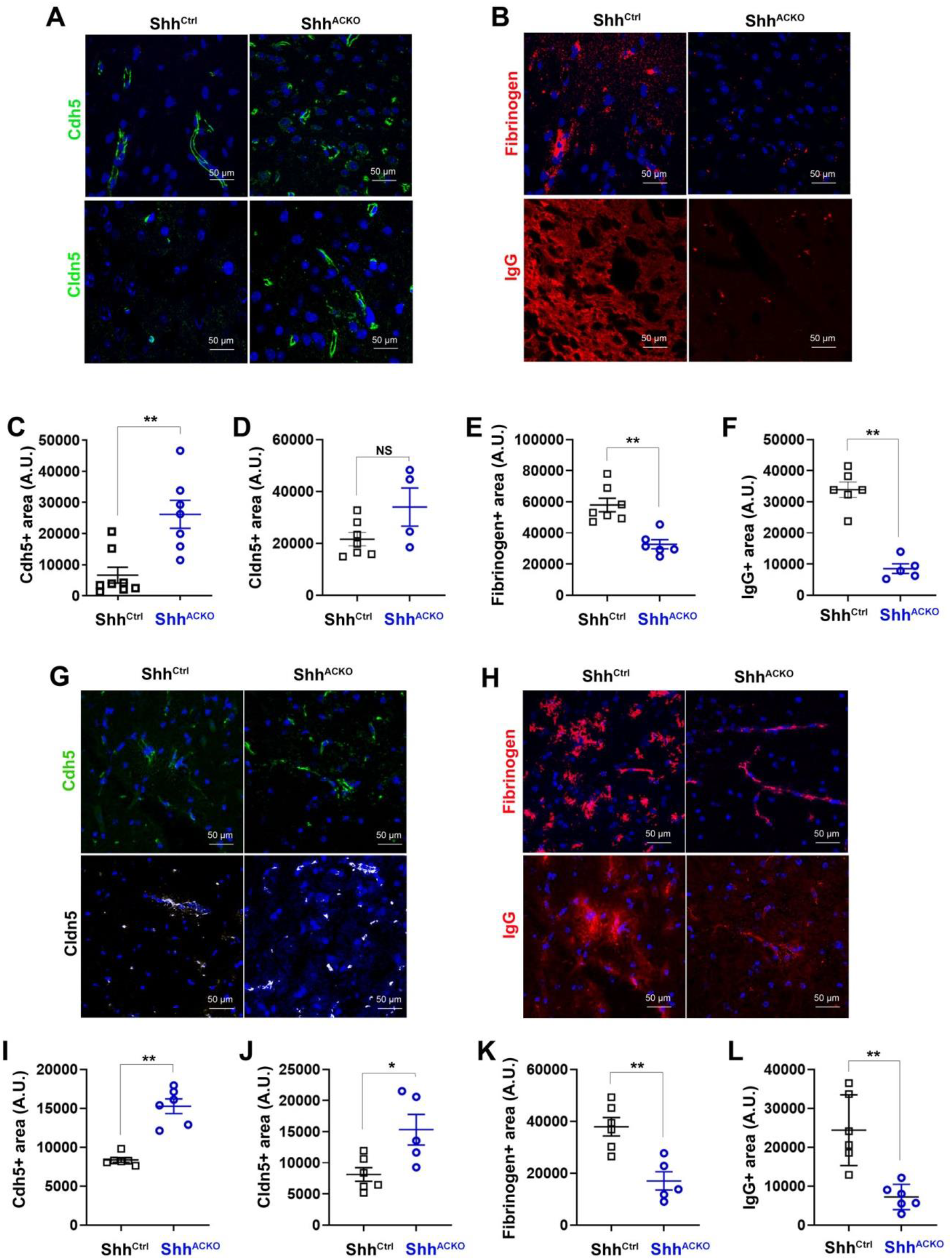
Astrocyte-derived Shh promotes BBB opening in the setting of CNS inflammation. (**A-F**) Both Glast-Cre^ERT2^; Shh^Flox/Flox^ (Shh^ACKO^) and Shh^Flox/Flox^ (Shh^ctrl^) mice were administered in the cerebral cortex with adenoviruses encoding Il1β (n=6 and 7 mice respectively). Mice were sacrificed 7 days later. (**A**) Brain sagittal sections were immunostained with anti-Cdh5 (in green) or anti-Cldn5 (in green) antibodies. Representative confocal images are shown. (**B**) Brain sagittal sections were immunostained with anti-Fibrinogen (in red) and anti-IgG (in red) antibodies. Representative confocal images are shown. Cdh5 (**C**) and Cldn5 (**D**) expression was quantified as the Cdh5+ and Cldn5+ surface area respectively. Fibrinogen (**E**) and Albumin (**F**) extravasation was quantified as the fibrinogen+ surface area and Albumin+ surface area respectively. (**G-L**) EAE was induced in both Shh^ACKO^ and Shh^ctrl^ mice (n=6 mice in each group). Mice were sacrificed 32 days later. (**G**) Spinal cord sections were immunostained with anti-Cdh5 (in green) and anti-Cldn5 (in white) antibodies. Representative confocal images are shown. (**H**) Spinal cord sections were immunostained with anti-Fibrinogen (in red) and anti-IgG (in red) antibodies. Representative confocal images are shown. Cdh5 (**I**) and Cldn5 (**J**) expression was quantified as the Cdh5+ and Cldn5+ surface area respectively. Fibrinogen (**K**) and IgG (**L**) extravasation was quantified as the Fibrinogen+ surface area and Albumin+ surface area respectively. *: p≤0.05; **: p≤0.01. Mann Withney test.

### Mice with astrocyte Shh inactivation display reduced disability in a model of multiple sclerosis

To examine the impact of these findings on disease severity, we investigated the phenotype of the multiple sclerosis model (EAE) in Shh^ACKO^ and control mice. Mice were sensitized with the encephalitogenic myelin peptide MOG_35–55_ and sacrificed 32 days later. Consistent with the acute cortex inflammation model, both Cdh5 and Cldn5 were significantly increased at the endothelium of spinal cord vessels in the absence of astrocyte-derived Shh (Figure 6F-H). Accordingly, fibrinogen and albumin extravasation was reduced (Figure 6F, I-J). These results thus confirmed the damaging role of astrocytic Shh at the BBB. As a consequence, in the absence of astrocyte-derived Shh, leucocyte entry in the spinal cord parenchyma was reduced (Figure 7A-B), so was microglia (Figure 7C-D) and astrocyte activation (Figure 7E-F) which all together concur to prevent demyelination (Figure 7G-H). Finally, the clinical consequences of astrocytic Shh deficiency were evaluated using a widely accepted 5-point paradigm from day 7 until the end of the experiment at day 32 after sensitization (Figure 7I). Critically, these studies revealed that the clinical course and pathology of EAE were strongly reduced in Shh^ACKO^ mice during the plateau of the disease. In controls, neurologic deficit was observed from day 9, and increased in severity until day 29, when clinical score stabilized at a mean of 2,91, representing hind limb paralysis. In contrast, the onset of clinical signs in Shh^ACKO^ mice was first seen 3 days later, and the clinical course was much milder. In Shh^ACKO^ mice, disease reached a plateau at day 19 at a mean of 1,78, indicating hind limb weakness and unsteady gait, a mild phenotype, and this divergence in scores are significant from day 8 to day 32 after sensitization (Figure 7I). The peak EAE score (1.75 vs 2.86 in control mice), total average score (0.86 vs 1.20 in control mice), and score during the time of disability (1.31 vs 1.82 in control mice) were all decreased in Shh^ACKO^ mice (data not shown) but there were no significant changes in survival and mortality rates (data not shown). In conclusion, this last set of data reveals for the first time that N-Shh may indeed act as a Dhh antagonist in a pathophysiological setting especially brain inflammation.

**Figure 7:**
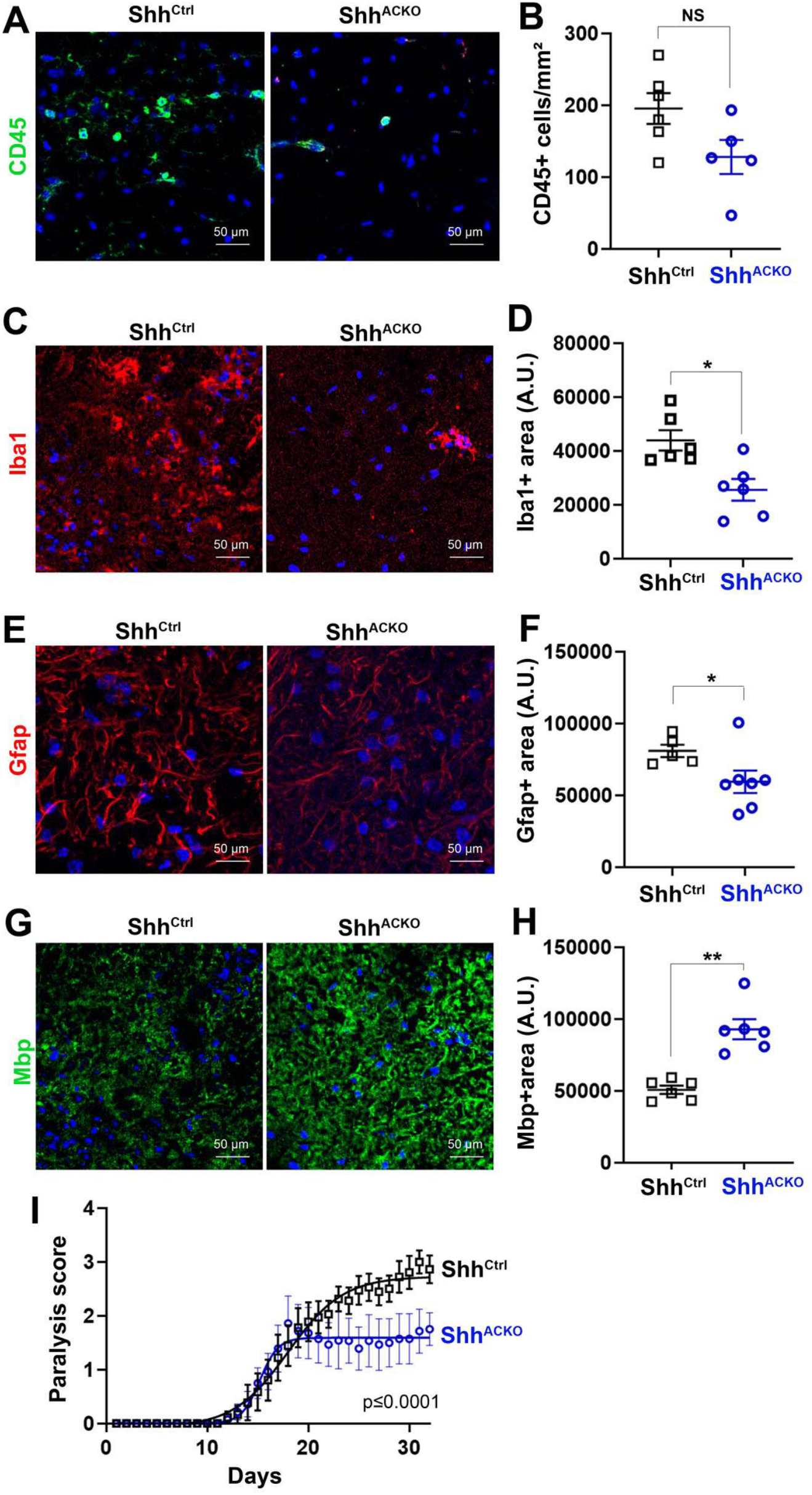
Astrocyte-derived Shh exacerbate EAE lesion formation and disease severity. EAE was induced in both Glast-Cre^ERT2^; Shh^Flox/Flox^ (Shh^ACKO^) and Shh^Flox/Flox^ (Shh^ctrl^) mice (n=6 mice in each group). (**A-H**) Mice were sacrificed 32 days later. (**A**) Spinal cord cross sections were immunostained with anti-CD45 antibodies to identify leucocytes. Representative confocal images are shown. (**B**) Leucocyte infiltration was quantified as the number of CD45+cells/mm². (**C**) Spinal cord cross sections were immunostained with anti-Iba1 antibodies to identify microglia. (**D**) Microglia activation was quantified as the Iba1+ surface area. (**E**) Activated Astrocytes were visualized after GFAP staining of Spinal cord cross sections. (**F**) Astrocyte activation was quantified as the Gfap+ surface area. (**G**) Spinal cord cross sections were immunostained with anti-Mbp antibodies to identify myelin. (**H**) Myelination was quantified as the MBP+ surface area. (**I**) Mice were scored daily from day 7 until the end of the experiment at day 32 after sensitization on a standard 5-point scale, nonlinear regression (Boltzmann sigmoidal). *: p≤0.05; **: p≤0.01; NS: not significant; Mann Withney test.

## Discussion

While Hh ligands are typically used instead of another for therapeutic purposes ^29^, the present study is the first one to our knowledge reporting that each forms of Hh ligand may induce its own specific effects. As a consequence, Hh ligands can enter into competition between each other to induce their effects. Notably, this is not the first time that different form of Hh ligand were reported to compete between each other, indeed, more than 20 years ago, unpalmitoylated Hh ligands were reported to act as dominant inhibitors toward palmitoylated Shh by competing for Ptch1 binding ^30^. These finding together with our finding thus highlight a totally new way of Hh signaling regulation based on the regulation of Hh ligand processing.

More specifically, in this study, we have investigated the effects of Hh ligands on ECs and compared the autocrine effects of 43 kDa FL-Dhh to the paracrine effects of 19 kDa N-Shh. We found that 43 kDa FL-Dhh promotes endothelial barrier integrity and inhibits angiogenesis while 19 kDa N-Shh destabilize endothelial intercellular junctions and promotes angiogenesis at least in part by acting as a competitive inhibitor of 43 kDa FL-Dhh demonstrating that Hh ligands cannot be used instead of another for therapeutic purposes. Moreover, 17 kDa N-Dhh also acts as a competitive inhibitor of 43 kDa FL-Dhh demonstrating that the different form of Hh ligands can also not be used instead of another.

While Hh canonical Hh signaling is typically induced in a “mesemchymal type” cell by a fully process 19 kDa N-Shh form produced by an “epithelial like” cells, Hh signaling in ECs occur through a non-canonical autocrine signaling. Consistent with the present finding such signaling was shown to be induced by full length-unprocessed Hh ligands ^31^. However the molecular mechanisms by which 43 kDa FL-Dhh promotes endothelial barrier function remains to be characterized. All that we know so far is that 43 kDa FL-Dhh-induced endothelial barrier integrity involves Smo but is more likely independent on Gli transcription factors. To further characterize this pathway is critical to fully understand why 43 kDa FL-Dhh, 45 kDa FL-Shh, 17 kDa N-Dhh and 19 N-Shh do not all induce the same effects in ECs. Indeed, while they all induce Ptch1 degradation through the proteasome, each of them induces a specific Ptch1 re-localization. Notably, Ptch1 inhibition and internalization were shown to be carried out by separable parts of Shh. ^32^ explaining why some Hh ligands-induced effects may be induced by any form of Hh ligands while other may be specific for one form or another. Consistently, it is currently not known why 19 N-Shh is a strong activator of Gli1 transcription in fibroblast while 17 N-Dhh is not ^22^.

Importantly, the present study may resolve quite a few inconsistent literature data regarding the role of Hh signaling in vascular biology ^14^. Indeed, in most studies that have investigated Shh effect on angiogenesis, Shh was delivered ectopically under its N-terminal form and was shown to be pro-angiogenic ^1,5,15^ which is fully consistent with the results obtained in the present study. However, it would be necessary to determine the cell origin and the form under which endogenous Shh is produced in the setting of ischemia to apprehend why ischemia-induced angiogenesis was shown to be accelerated in Shh deficient mice ^16^. Besides, studies reporting that Hh signaling promotes BBB integrity in adults have either used Smo inhibitors, Smo^ECKO^ or Dhh^ECKO^ mice ^2,17,33^. In each of these cases, 43 kDa FL-Dhh-induced autocrine signaling in ECs was then more likely responsible for the findings. On the contrary, when glioblastoma-derived Dhh was shown to disrupt the BBB ^18^, it may have been produced under its 17 kDa N-Dhh form which compete with 43 kDa FL-Dhh. The same conclusion can be made from the study reporting that ectopic Shh expression induces hemorrhage in the spinal cord ^19^. Once again either FL-Shh or N-Shh may have act as 43 kDa FL-Dhh competitive antagonist.

Moreover, while Hh signaling is mostly shown to be good for the cardiovascular system, Scube2 was shown to be upregulated in the setting of type 2 diabetes ^34^ and atherosclerosis ^35^. Besides Scube2 has been associated with endothelial dysfunction ^34^ and has a strong proangiogenic activity ^36,37^. Since Scube2 promotes soluble 19 kDa cholesterolated N-Shh releases. These results are actually fully consistent with the conclusion of the present study i.e. 43 kDa FL-Dhh promotes endothelial barrier integrity and inhibits angiogenesis while 19 kDa N-Shh destabilize endothelial intercellular junctions and promotes angiogenesis.

In conclusion, the present study reveals unsuspected competitive and antagonistic activities of the different forms and types of Hh ligands in the adult cardiovascular systems especially ECs. Such results represent a breakthrough regarding our understanding of the regulation of Hh signaling and demonstrate that Hh ligands or form of Hh ligands cannot be used instead of another for therapeutic purposes.

## Methods

### Mice

Dhh Floxed (Dhh^Flox^) mice were generated at the “Institut Clinique de la Souris” through the International Mouse Phenotyping Consortium (IMPC) from a vector generated by the European conditional mice mutagenesis program, EUCOMM and described before ^17^. Shh Floxed (Shh^Flox^) mice ^38^, Smo Floxed (Smo^Flox^) mice ^39^, Tg(Slc1a3-cre/ERT)1Nat/J (Gast-Cre^ERT2^) mice ^40(p2)^ and *Gt(ROSA)26Sor^tm4(ACTB-tdTomato,-EGFP)Luo^*/J (Rosa26^mTmG^) mice ^41^ were obtained from the Jackson Laboratories. Tg(Cdh5-cre/ERT2)1Rha (Cdh5-CreERT2) mice ^42^ were a gift from RH. Adams.

Animal experiments were performed in accordance with the guidelines from Directive 2010/63/EU of the European Parliament on the protection of animals used for scientific purposes and approved by the local Animal Care and Use Committee of Bordeaux University.

The Cre recombinase in Cdh5-Cre^ERT2^ mice was activated by intraperitoneal injection of 1 mg tamoxifen for 5 consecutive days at 8 weeks of age. Mice were phenotyped 2 weeks later. Successful and specific activation of the Cre recombinase has been verified before ^43^. Successful and specific activation of the Cre recombinase has been verified before ^43^. The Cre recombinase in Gast-Cre^ERT2^ mice was activated by intraperitoneal injection of 1 mg tamoxifen, twice a day for 5 consecutive days at 8 weeks of age. Mice were phenotyped 2 weeks later. Successful and specific activation of the Cre recombinase has been verified (Supplementary Figure 7A).

Both males and females were used in equal proportions except otherwise stated.

### Mouse corneal angiogenesis assay

Pellets were prepared as previously described ^44^. Briefly, either 5 μg of VEGFA (Shenandoah biotechnology) or rec N-Shh (Shenandoah biotechnology) diluted in 10 μL sterile phosphate-buffered saline (PBS) was mixed with 2.5 mg sucrose octasulfate-aluminum complex (Sigma-Aldrich Co., St. Louis, MO, USA), and 10 μL of 12% hydron in ethanol was added. The suspension was deposited on a 400-μm nylon mesh (Sefar America Inc., Depew, NY, USA), and both sides of the mesh were covered with a thin layer of hydron and allowed to dry.

Female mice were anesthetized with an intraperitoneal (IP) injection of ketamine 100 mg/kg and xylazine 10 mg/kg. The eyes of the mice eyes were topically anesthetized with 0.5% Proparacaine™ or similar ophthalmic anesthetic. The globe of the eye was proptosed with jeweler’s forceps taking care to not damage the limbus vessel surrounding the base of the globe. Sterile saline was also be applied directly to each eye as needed during the procedure to prevent excessive drying of the cornea and to facilitate insertion of the pellet into the lamellar pocket of the eyes. Using an operating microscope, a central, intrasomal linear keratotomy was performed with a surgical blade parallel to the insertion of the lateral rectus muscle. Using a modified von Greafe knife, a lamellar micro pocket was made toward the temporal limbus by ‘rocking’ the von Greafe knife back and forth. The pellet was placed on the cornea surface with jeweler’s forceps at the opening of the lamellar pocket. A drop of saline was applied directly to the pellet, and using the modified von Greafe knife, the pellet was gently advanced to the temporal end of the pocket. Buprenorphine was given at a dose of 0.05 mg/kg subcutaneously on the day of surgery.

Nine days after pellet implantation, mice were sacrificed, and eyes were harvested and fixed with 2% formalin. Capillaries were stained with rat anti-mouse CD31 antibodies (BMA Biomedicals, Cat#T-2001), and primary antibodies were visualized with Alexa 568–conjugated anti-rat antibodies (Invitrogen). Pictures were taken under 50x magnification. Angiogenesis was quantified as the CD31+ surface area using Image J software.

### Central nervous system microinjection

To obtain LV-N-Shh, pIRES-N-Shh ^16^ was digested by *NheI* and *HpaI*, the mouse N-Shh sequence was then cloned in pRRLsin.MND.MCS.WPRE (Addgene) that had been previously digested by *NheI* and *PmlI*. Lentviral particles were produced at the vectorology facility of Bordeaux University.

Mice were anaesthetized using isoflurane and placed into a stereotactic frame (Stoelting, Illinois, USA). To prevent eye dryness, an ophthalmic ointment was applied at the ocular surface to maintain eye hydration during the time of surgery. The skull was shaved and the skin incised on 1 cm to expose the skull cap. Then, a hole was drilled into the cerebral cortex and 3 μL of an AdIL-1 ^45^, AdDL70 control (AdCtrl), LV-NShh or LV-GFP control (10^7^ pfu) solution was microinjected at y=1 mm caudal to Bregma, x=2 mm, z=1.5 mm using a Hamilton syringe, into the cerebral cortex and infused for 3 minutes before removing the needle from the skull hole ^46^. Mice received a subcutaneous injection of buprenorphine (0.05 mg/kg) 30 minutes before surgery and again 8 hours post-surgery to assure a constant analgesia during the procedure and postoperatively. Mice were sacrificed by pentobarbital overdose. Those injected with the adenovirus were sacrificed 7 days post-surgery and those injected with the lentivirus were sacrificed 14 days post-surgery to maximize the vector effectiveness. For histological assessment, brains were harvested and fixed in formalin for 3 hours before being incubated in 30% sucrose overnight and OCT embedded. Then, for each brain, the lesion area identified by the puncture site was cut into 7 μm thick sections.

### Experimental autoimmune encephalomyelitis (EAE)

Ten week old female mice were immunized by subcutaneous injection of 300 μg myelin oligodendrocyte glycoprotein-35-55 (MOG35–55) (Hooke laboratories) in 200-μl Freund’s adjuvant containing 300 μg/mL mycobacterium tuberculosis H37Ra (Hooke laboratories) in the dorsum. Mice were administered with 500 ng pertussis toxin (PTX) IP on day of sensitization and 1 day later (Hooke laboratories). Disease was scored (0, no symptoms; 1, floppy tail; 2, hind limb weakness (paraparesis); 3, hind limb paralysis (paraplegia); 4, fore- and hind limb paralysis; 5, death) ^45^ from day 7 post immunization until day 32 post immunization. At day 32, all the animals were euthanized by pentobarbital overdose. For histological assessment, cervical, lumbar and dorsal sections of each animal spinal cord, as well as the spleen, were harvested and fixed in formalin for 3 hours before being incubated in 30% sucrose overnight, embedded in OCT and cut into 9 μm thick sections.

### Immunostaining

Prior to staining, brains were fixed in 10% formalin for 3 hours, incubated in 30% sucrose overnight, OCT embedded and cut into 7 μm thick sections. Cultured cells were fixed with 10% formaline for 10 minutes.

Human Cdh5 was stained using mouse anti-human Cdh5 antibodies (Santa Cruz Biotechnology, Inc, Cat#sc-9989). Mouse Cdh5 was stained using goat anti-mouse Cdh5 antibodies (R&D systems, Cat# AF1002). Cldn5 was stained using either mouse monoclonal anti-Cldn5 antibodies (Invitrogen, Cat# 35-2500) or rabbit polyclonal anti-Cldn5 antibodies (Invitrogen, Cat# 34-1600). Albumin and fibrinogen were stained using sheep anti-albumin antibodies (Abcam, Cat# ab8940) and rabbit anti-fibrinogen antibodies (Dako, Cat#A0080) respectively.

Pan-leucocytes were identified using rat anti-mouse CD45 antibodies (BD Pharmingen Inc, Cat# 550539). Iba1+ microglia was identified using rabbit anti-Iba1 antibodies (FUJUFILM Wako chemicals, cat#019-19741). Reactive astrocytes were stained using rabbit anti-Gfap antibodies (ThermoFisher, Cat# OPA1-06100). Myelin was identified using rat anti-Mbp antibodies (Abcam, Cat# ab7349). Hh ligands were stained using rabbit anti-Shh antibodies (Santa Cruz Biotechnology, Inc, Cat# sc-9024). Dhh was specifically stained using mouse anti-Dhh antibodies (Santa Cruz Biotechnology, Inc, Cat# sc-271168). Ptch1 was stained using rabbit anti-Ptch1 antibodies (Abcam, Cat# ab53715). For immunofluorescence analyzes, primary antibodies were resolved with Alexa Fluor^®^–conjugated secondary polyclonal antibodies (Invitrogen, Cat# A-21206, A-21208, A-11077, A-11057, A-31573, A-10037) and nuclei were counterstained with DAPI (1/5000). For both immunohistochemical and immunofluorescence analyses, negative controls using secondary antibodies only were done to check for antibody specificity.

BBB permeability was evaluated by measuring tight junction integrity and plasmatic protein extravasation. For each brain or spinal cord section, Cldn5+, Cdh5+, Fibrinogen+, and Albumin+ and IgG+ areas were quantified in 20 pictures taken at the margins of the lesion area under 40x magnification. One section was quantified per brain (the section is localized in the Il1β-induced inflammatory lesion area) or spinal cord (for the spinal cord, 3 different zones are displayed on the same section: one cervical, one lumbar and one dorsal to get a global vision of the lesion) (per mouse).

For each spinal cord section, CD45+ leukocytes were counted in 20 pictures randomly taken under 40x magnification. Three sections/per mice was quantified per spinal cord (one cervical, one lumbar and one dorsal) to get a global vision of the inflammatory lesion). For each spinal cord section, Gfap+ and NeuN+ areas were quantified in 10 pictures taken in and around the lesion area under 20X magnification. Three sections/per mice was quantified per spinal cord (one cervical, one lumbar and one dorsal).

### Isolation of primary cultured mouse brain ECs

Mice were sacrificed by cervical dislocation. Brain was then harvested and cerebellum, olfactory bulb and white matter removed with sterile forceps. Additionally, meninges were eliminated by rolling a sterile cotton swab at the surface of the cortex. The cortex was then transferred in a potter containing 2 mL HBSS 1X w/o phenol red containing 10 mol/L HEPES and 0,1% BSA and the brain tissue was pounded to obtain an homogenate which was collected in a 15 mL tube. Cold 30% dextran solution was then added to the tube (V/V) to obtain a 15% dextran working solution. After a 25 min centrifugation at 3000 g at 4°C, the pellet (neurovascular components and red cells) was collected and the supernatant (dextran solution and neural components) was centrifuged again to get the residual vessels. Neurovascular components were then pooled and washed three time with HBSS 1XCa^2+^/Mg^2+^ free containing phenol red, 10 mmol/LHEPES and 0,1% BSA. The pellet was then suspended in DMEM containing 2 mg/mL collagenase/dispase (Roche), 0,147 μg/mL TLCK (Lonza) and 10 μg/mL DNAse 1 (Roche), pre-warmed at 37 °C, before being placed on a shaking table at maximum speed agitation at 37°C. After 30 min, the digestion was stopped by adding 10 mL HBSS 1X Ca^2+^/Mg^2+^ free containing phenol red, 10 mmol/L HEPES and 0,1% BSA and washed again 3 times in this same buffer. The digested neurovascular pellet was finally re-suspended in mouse brain endothelial cell culture medium (DMEM 1 g/L glucose containing 20% FBS, 2% sodium pyruvate, 2% non-essential amino acids, 1 ng/mL FGF2 and 10 mg/mL gentamycin) and plated on dishes previously coated with 2% Matrigel™ (BD biosciences).

### Cell culture

Human umbilical vein endothelial cells (HUVECs) (Lonza) were cultured in endothelial basal medium-2 (EBM-2) supplemented with EGM™-2 BulletKit™ (Lonza). Cell from passage 3 to passage 6 were used. HeLa ATCC^®^CCL-2™ cells were cultured in Roswell Park Memorial Institute medium (RPMI) supplemented with 10% fetal bovine serum. Normal human astrocytes (Lonza) were cultured in astocytic basal medium (ABM) supplemented with AGM bulletKit™ (Lonza). Cell from passage 2 to passage 5 were used. EA.hy926 ATCC ^®^ CRL-2922™ were cultured in DMEM containing 1 g/L glucose and supplemented with 10% fetal bovine serum.

### siRNA/transfection

HUVECs were transfected with human Dhh siRNA: rArCrUrCrCrUrUrArArArGgArGrGrArCrUrArUrUrUrArGCC, human Disp1 siRNA: rGrGrArUrCrUrArArCrArArGrUrUrArCrArUrUrGrUrArUAG or universal scrambled negative control siRNA duplex (Origen) using JetPRIME™ transfection reagent (Polyplus Transfection), according to the manufacturer’s instructions.

### Plasmids/Transfection

pIRES-NDhh ^47^, pIRES-NShh ^16^ and pcDNA3-FL-Dhh ^43^ were described previously. Human FL-Dhh sequence was obtained after EcoRI digestion of pBS hSHH (Addgene ID13996) and then cloned at the EcoRI site of pCDNA3.1 myc His to generate the pcDNA3-FL-Shh plasmid. The myc-tagged human Ptch1, Ptch1-1B-myc was kindly given by R. Toftgard ^48^ and the human full length Dhh was previously described ^43^. HeLa cells were transfected using JetPRIME™ transfection reagent (Polyplus Transfection), according to the manufacturer’s instructions.

### Quantitative RT-PCR

RNA was isolated using Tri Reagent^®^ (Molecular Research Center Inc) as instructed by the manufacturer, from 3×10^5^ cells or from tissue that had been snap-frozen in liquid nitrogen and homogenized. For quantitative RT-PCR analyses, total RNA was reverse transcribed with M-MLV reverse transcriptase (Promega) and amplification was performed on a DNA Engine Opticon^®^2 (MJ Research Inc) using B-R SYBER^®^ Green SuperMix (Quanta Biosciences). Primer sequences are reported in Supplementary table 1.

The relative expression of each mRNA was calculated by the comparative threshold cycle method and normalized to Actb mRNA expression.

### Migration assay

Cell migration was evaluated with a chemotaxis chamber (Neuro Probe, Inc., Gaithersburg, MD, USA). Briefly, a polycarbonate filter (8 μm pore size) (GE Infrastructure, Fairfield, CN, USA) was coated with a solution containing 0.2% gelatin (Sigma-Aldrich Co.) and inserted between the chambers. 5×10^4^ cells per well were seeded in the upper chamber, and the lower chamber was filled with EBM-2 containing 0.5% FBS. Cells were incubated for 8 hours at 37°C then viewed under 20× magnification, and the number of cells that had migrated to the lower chamber were counted in 3 HPFs per well; migration was reported as the mean number of migrated cells per HPF. Each condition was assayed in triplicate and each experiment was performed at least three times.

### In vitro permeability assay

100 000 cells were seeded in Transwell^®^ inserts. The day after, 0.5 mg/mL 70 kDa FITC-dextran (Sigma) was added to the upper chamber. FITC fluorescence in the lower chamber was measured one hour later.

### Immunoprecipitation/western blot analysis

Prior to western blot analysis, Ptch1 was immunoprecipitated with anti myc-tag antibodies (Millipore, Cat# 05-724). Expression of Dhh, Shh, Ptch1 and Smo were evaluated by SDS PAGE using mouse anti-Dhh antibodies (Santa Cruz Biotechnology, Inc, Cat# sc-271168), mouse anti-Shh antibodies (Santa Cruz Biotechnology, Inc, Cat#sc-365112), rabbit-anti Hh antibodies (Santa Cruz Biotechnology, Inc, Cat# sc-9024), rabbit anti-Ptch1 antibodies (Abcam, Cat#ab53715).

Expression of junction protein Cdh5, Ocln and Cldn5 was evaluated by SDS PAGE using goat anti-mouse Cdh5 antibodies (R&D systems, cat#AF1002), mouse anti-Ocln (Invitrogen, cat#33-1500) and mouse anti-Cldn5 antibodies (Invitrogen, cat#33-1500.

Protein loading quantity was controlled using rabbit monoclonal anti-β-actin antibodies (Cell Signaling Technology, Cat#4970). Secondary antibodies were from Invitrogen, Cat#A-21039, A-21084, A-21036). The signal was then revealed by using an Odyssey Infrared imager (LI-COR).

### Proximity ligation assay

Proximity ligation assay was performed using the Duolink^®^ In Situ Orange Starter Kit Mouse/Rabbit (Sigma) according to the manufacturer instructions. Images were taken at the 63x magnification under a confocal microscope. 8-12 transfected cells per conditions were imaged for quantification.

### Statistics

Results are reported as mean ± SEM. Comparisons between groups were analyzed for significance with the non-parametric Mann-Whitney test or a Kruskal-Wallis test followed by Dunn’s multiple comparison test (for than two groups) using GraphPad Prism v8.0.2 (GraphPad Inc, San Diego, Calif). Differences between groups were considered significant when p≤0.05 (*: p≤0.05; **: p≤0.01; ***: p≤0.001).

## Author contributions

P.-L. H. and C.C. conducted experiments, acquired data, analyzed data. S.G. and L.C. conducted experiments, acquired data. A.-P. G. critically revised the manuscript. M.-A. R. designed research studies, conducted experiments, acquired data, analyzed data, providing reagents, and wrote the manuscript

## Acknowledgments

We thank Annabel Reynaud, Sylvain Grolleau, and Maxime David for their technical help. We thank Christelle Boullé for administrative assistance.

This study was supported by grants from the Fondation de France (Appel d’Offre Recherche sur les maladies Cardiovasculaires 2013 and 2018), and the Fondation ARSEP pour la recherche sur la sclérose en plaques. Also this study was funded by a Marie Skłodowska-Curie Actions (MSCA-IF-2019) from the European concil. Finally, this study was co-funded by the “Institut National de la Santé et de la Recherche Médicale” and by the University of Bordeaux.

