## Supplementary data for "Endothelial cell response to Hedgehog ligands depends on their processing"

**Supplementary table**

| | | |
|---|---|---|
| hActb | **F** | 5′-GGAGGAGCTGGAAGCAGCC-3′ |
| | R | 5′-GCTGTGCTACGTCGCCCTG-3′ |
| hGli1 | F | 5′-TTCCTACCAGAGTCCCAAGT-3′ |
| | R | 5′-CCCTATGTGAAGCCCTATTT-3′ |
| hGli2 | F | 5′-CAGATCCACATGTACGAACAG-3′ |
| | R | 5′-CCATGATGGCATCGAAGTC-3′ |
| hShh | F | 5′-CAGTTTATCCCCAATGTGGC-3′ |
| | R | 5′-GCCAAAGCGTTCAACTTGTC-3′ |
| mActb | F | 5′-CCTGAACCCTAAGGCCAACC-3′ |
| | R | 5′-TAGCCCTCGTAGATGGGCAC-3′ |
| mGli1 | F | 5′-GAAGGAATTCGTGTGCCATT-3′ |
| | R | 5′-GCAACCTTCTTGCTCACACA-3′ |
| mGli2 | F | 5′-TGGTTGAGCGGAAGGTTGAA-3′ |
| | R | 5′-ACAGGTTTGGGATACGCACA-3′ |

F forward, R reverse
Actb was used as the household gene

**Supplementary Table I: List of primers used for reverse transcription (RT) quantitative polymer chain reaction (qPCR)**

### Supplementary Figures and Supplementary Figure legends

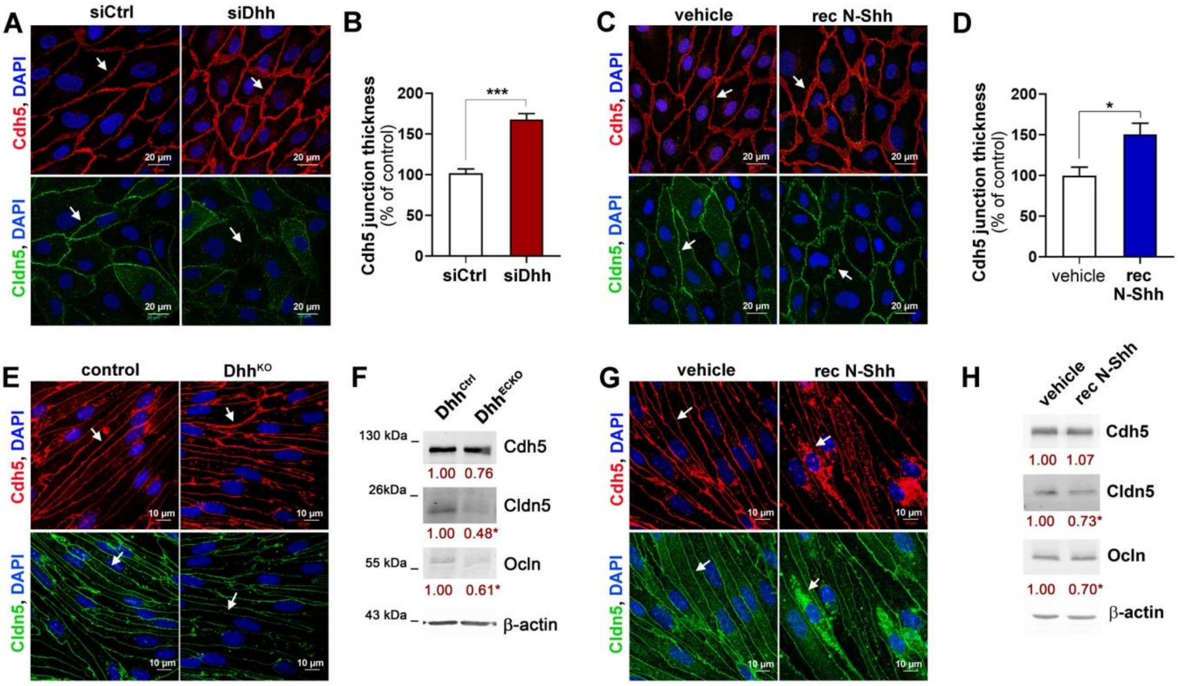


**Supplementary Figure 1:** ***EC-derived Dhh improves intercellular junction integrity while recombinant N-Shh disrupts it.*** (**A-B**) HUVECs were transfected with Dhh or control siRNAs. (**A**) Cdh5 (in red) and Cldn5 (in green) localization was evaluated by immunofluorescent staining of a confluent cell monolayer and (**B**) quantified as the mean junction thickness using Image J software. The experiment was repeated at least 4 times. (**C-D**) HUVECs were treated or not with 1 µg/mL recombinant N-Shh (rec N-Shh) for 30 minutes. (**C**) Cdh5 (in red) and Cldn5 (in green) localization was evaluated by immunofluorescent staining of a confluent cell monolayer and (**B**) quantified as the mean junction thickness using Image J software. The experiment was repeated at least 4 times. (**E-F**) mouse brain ECs were isolated from Cdh5-Cre$^{ERT2}$ Dhh$^{Flox/Flox}$ (Dhh$^{ECKO}$) and Dhh$^{Flox/Flox}$ (Dhh$^{Ctrl}$) mice that had been previously administered with tamoxifen (n=4 in each group).(**E**) Cdh5 (in red) and Cldn5 (in green) localization was evaluated by immunofluorescent staining of a confluent cell monolayer. (**F**) Cdh5, Cldn5 and Ocln protein levels were quantified by western blot analyses. (**G-H**) mouse brain ECs were isolated from WT mice, and then treated or not with 1 µg/mL rec N-Shh for 30 minutes. (**G**) Cdh5 (in red) and Cldn5 (in green) localization was evaluated by immunofluorescent staining of a confluent cell monolayer. (**H**) Cdh5, Cldn5 and Ocln protein levels were quantified by western blot analyses. *: p≤0.05; ***: p≤0.001. Mann Whitney test.

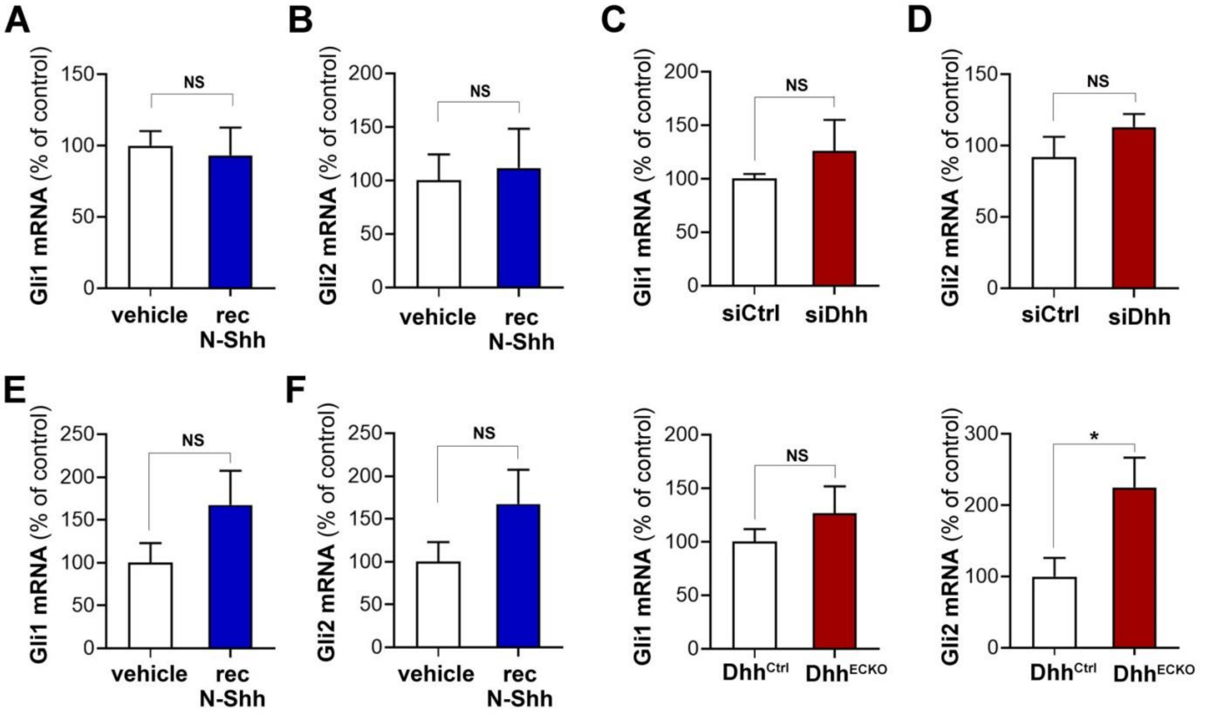


**Supplementary Figure 2:** ***Hh ligands do not modulate Gli expression in ECs*****.** (**A-B**) HUVECs were treated or not with 1 µg/mL recombinant N-Shh (recN-Shh) for 24 hours. Gli1 (**A**) and Gli2 (**B**) mRNA expression was quantified via RT-qPCR. The experiment was repeated 3 times, each experiment included triplicates. (**C-D**) HUVECs were transfected with Dhh or control siRNAs. Gli1 (**C**) and Gli2 (**D**) mRNA expression was quantified via RT-qPCR. The experiment was repeated 3 times, each experiment included triplicates. (**E-F**) Mouse brain ECs were isolated from WT mice, and then treated or not with 1 µg/mL rec N-Shh for 24 hours. Gli1 (**E**) and Gli2 (**F**) mRNA expression was quantified via RT-qPCR. The experiment was repeated 3 times, each experiment included triplicates. (**G-H**) mouse brain ECs were isolated from Cdh5-Cre$^{ERT2}$ Dhh$^{Flox/Flox}$ (Dhh$^{ECKO}$) and Dhh$^{Flox/Flox}$ (Dhh$^{Ctrl}$) mice that had been previously administered with tamoxifen (n=4 in each group). Gli1 (**G**) and Gli2 (**H**) mRNA expression was quantified via RT-qPCR. The experiment was repeated 3 times, each experiment included triplicates. NS: not significant. Mann Whitney test

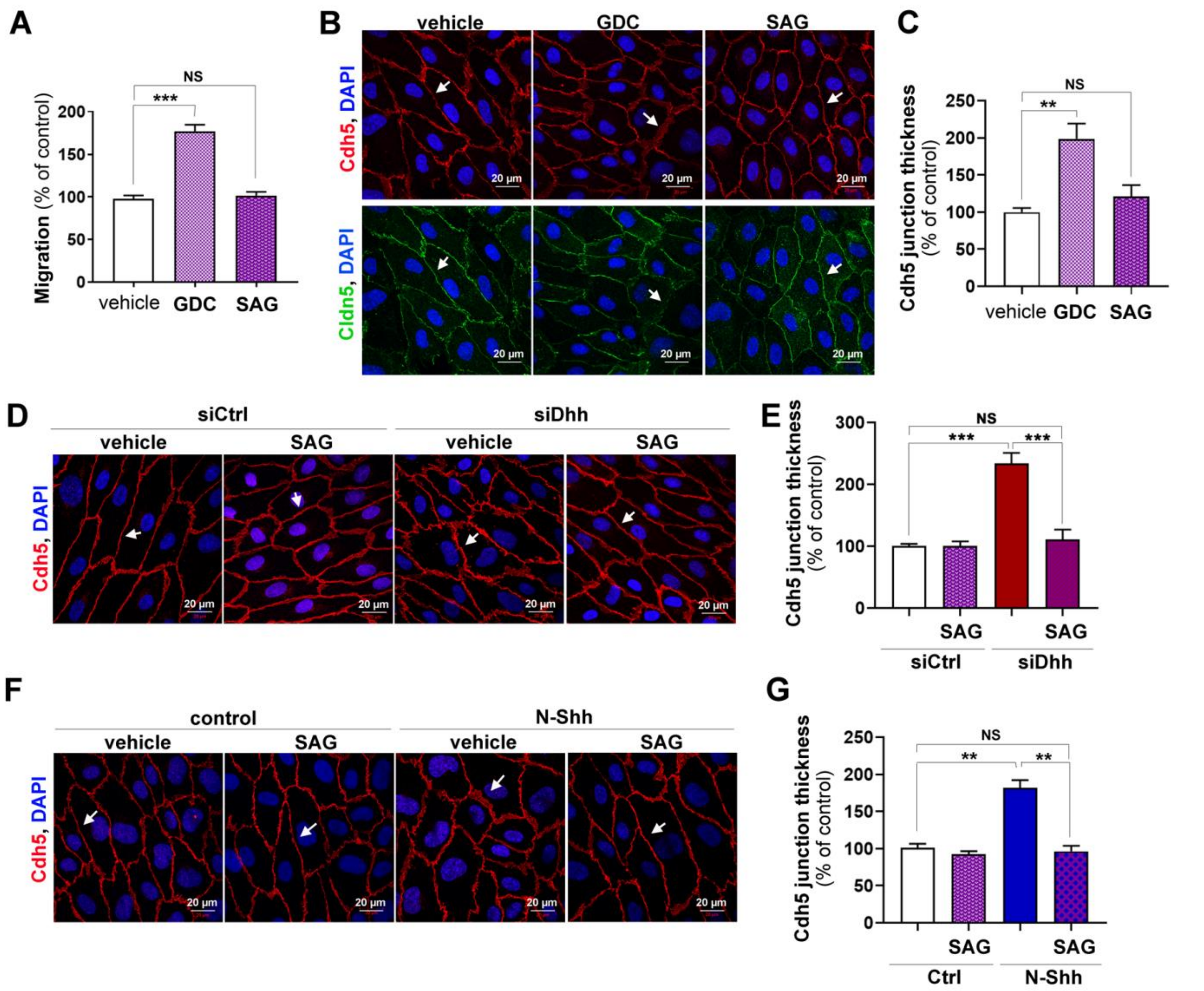


**Supplementary Figure 3:** ***Inhibition of Hh signaling in ECs destabilizes intercellular junctions sand promotes EC migration.*** (**A-C**) HUVECs were treated either with 30 nmol/L GDC-0449, 100 nmol/L SAG or control vehicle. (**A**) Cell migration was assessed in a chemotaxis chamber. The experiment was repeated 3 times, each experiment included n=4 wells/conditions. (**B**) Cdh5 (in red) and Cldn5 (in green) localization was evaluated by immunofluorescent staining of a confluent cell monolayer and (**C**) quantified as the mean junction thickness using Image J software. The experiment was repeated at least 4 times. (**D-E**) HUVECs were transfected with Dhh or control siRNAs and then treated or not with 100 nmol/L SAG for 16 hours (**D**) Cdh5 (in red) localization was evaluated by immunofluorescent staining of a confluent cell monolayer and (**E**) quantified as the mean junction thickness using Image J software. The experiment was repeated at least 4 times. (**F-G**) HUVECs were treated or not with 1 µg/mL rec N-Shh for 16 hours in the presence or not of 100 nmol/L SAG (**F**) Cdh5 (in red) localization was evaluated by immunofluorescent staining of a confluent cell monolayer and (**G**) quantified as the mean junction thickness using Image J software. The experiment was repeated at least 4 times.*: $p \leq 0.05$; **: $p \leq 0.01$; ***: $p \leq 0.001$; NS: not significant. Kruskal-Wallis test followed by Dunn's multiple comparison test.

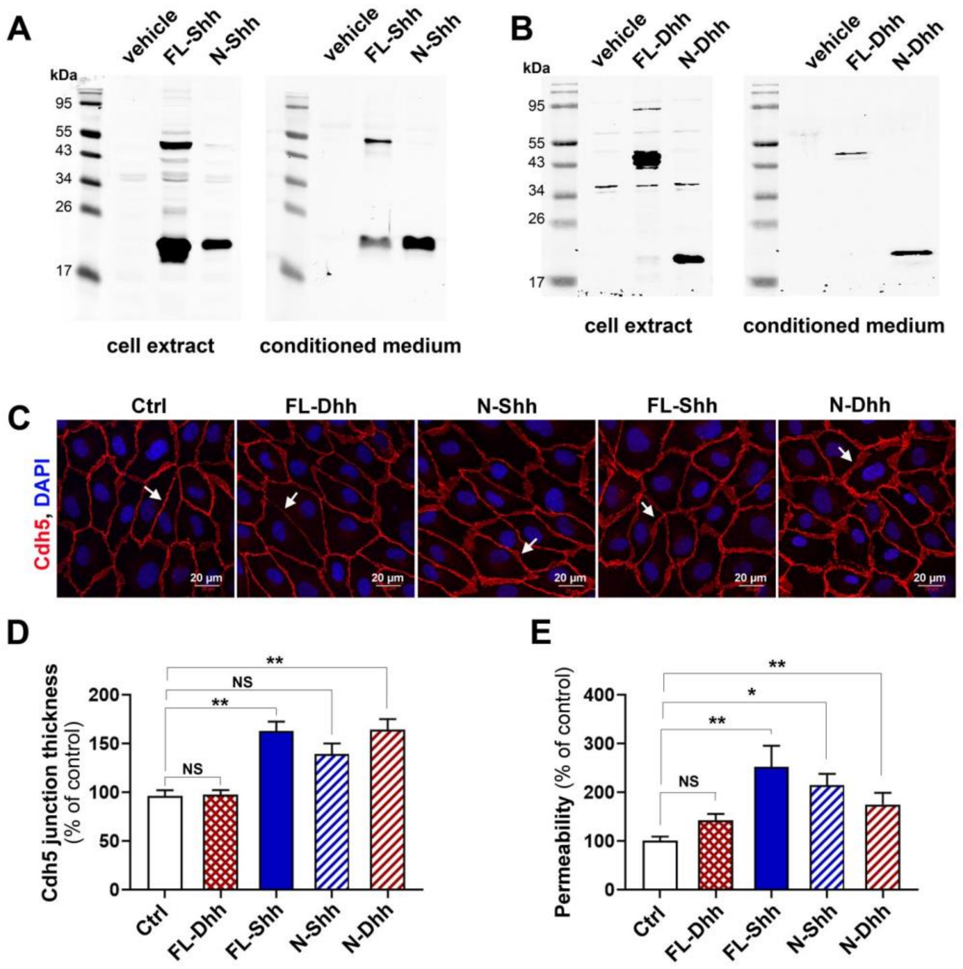


**Supplementary Figure 4:** (**A**) HeLa cells were transfected either with FL-Shh or N-Shh encoding plasmids. Expression and secretion of Shh was assessed by western blot analysis. (**B**) HeLa cells were transfected either with FL-Dhh or N-Dhh encoding plasmids. Expression and secretion of Dhh was assessed by western blot analysis. (**C-E**) HUVECs were treated with conditioned medium from HeLa containing or not FL-Dhh, N-Shh, FL-Shh or N-Dhh. (**C**) Cdh5 localization was evaluated by immunofluorescent staining (in red) of a confluent cell monolayer and (**D**) quantified as the mean junction thickness using Image J software. The experiment was repeated at least 4 times. (**E**) Endothelial monolayer permeability to 70 kDa FITC-Dextran was assessed using Transwells. The experiment was repeated 3 times, each experiment included triplicates. *: $p \leq 0.05$; **: $p \leq 0.01$; ***: $p \leq 0.001$; NS: not significant. Kruskal-Wallis test followed by Dunn's multiple comparison test.

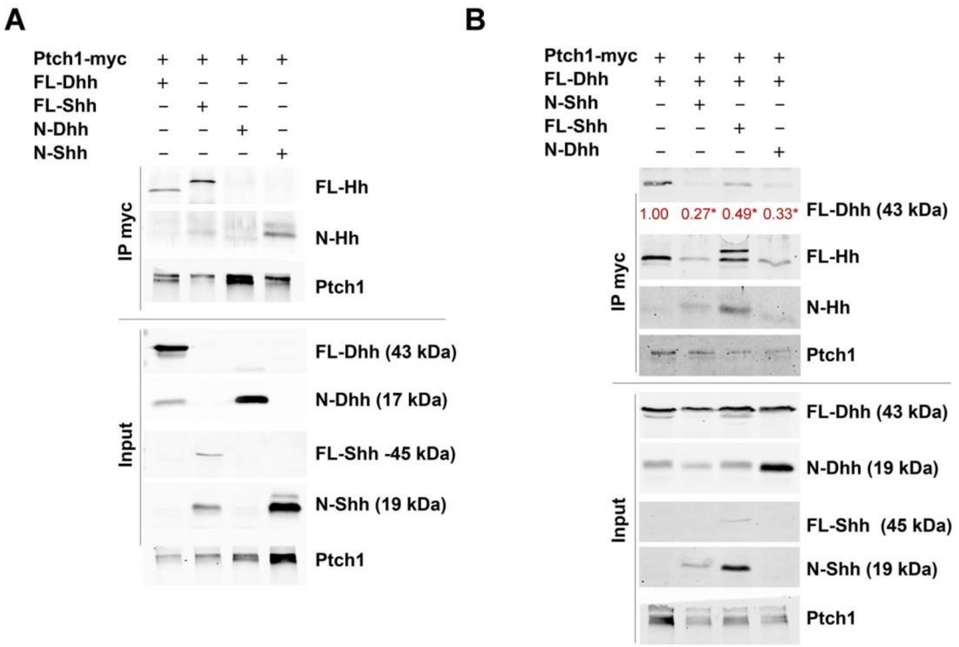


**Supplementary Figure 5:** (**A**) HeLa cells were co-transfected Ptch1-myc together with FL-Dhh, FL-Shh, N-Dhh or N-Shh encoding plasmids. Hh ligands interaction with Ptch1 was assessed by co-immunoprecipitation assay. (**B**) HeLa were co-transfected Ptch1-myc and FL-Dhh encoding plasmids together with or without N-Shh, FL-Shh or N-Dhh encoding plasmids. Hh ligands interaction with Ptch1 was assessed by co-immunoprecipitation assay. *: p≤0.05. Kruskal-Wallis test followed by Dunn's multiple comparison test.

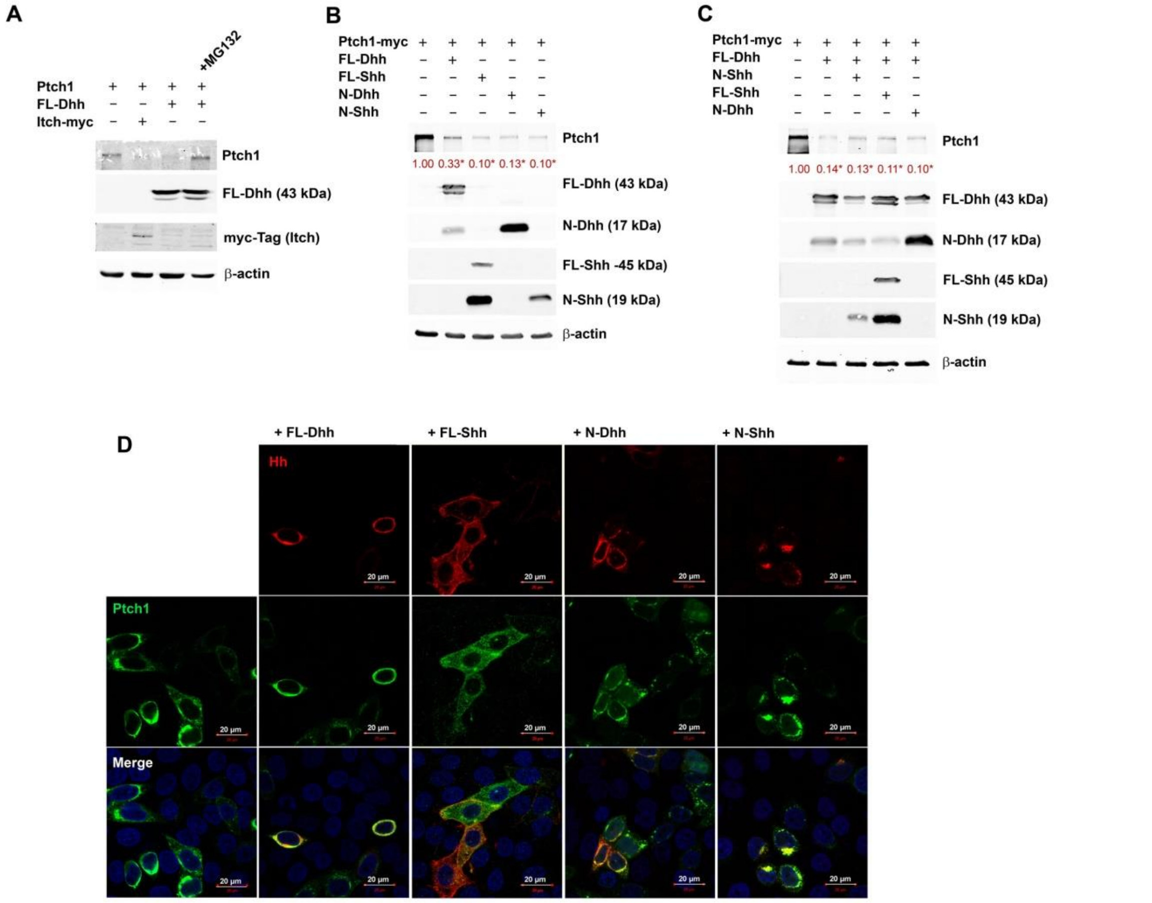


**Supplementary Figure 6:** (**A**) HeLa cells were co-transfected with Ptch1-myc together with Itch-myc or FL-Dhh encoding plasmids then treated or not with 10 µmol/L MG132 for 16 hours. Ptch1, Dhh, and Itch protein level was assessed by western blot analysis. (**B**) HeLa cells were co-transfected with Ptch1-myc encoding plasmids together with plasmids expressing or not FL-Dhh, FL-Shh, N-Dhh or N-Shh. Ptch1, Dhh, and Shh protein level was assessed by western blot analysis. The experiment was performed at least 4 times. (**C**) HeLa cells were co-transfected Ptch1-myc and FL-Dhh encoding plasmids together with or without N-Shh, FL-Shh or N-Dhh encoding plasmids. Ptch1, Dhh, and Shh protein level was assessed by western blot analysis. The experiment was performed at least 4 times. (**D**) HeLa cells were co-transfected with Ptch1-myc encoding plasmids together with plasmids expressing or not FL-Dhh, FL-Shh, N-Dhh or rec N-Shh. Ptch1 (in green), Dhh (in red), and Shh (in red) protein localization was assessed by immune-fluorescent staining. (**D**) EA.hy926 cells were co-transfected with Ptch1-myc encoding plasmids together with plasmids expressing or not FL-Dhh, FL-Shh, N-Dhh or N-Shh. Ptch1 (in green), Dhh (in red), and Shh (in red) protein localization was assessed by immune-fluorescent staining. *: p≤0.05. Kruskal-Wallis test followed by Dunn's multiple comparison test.

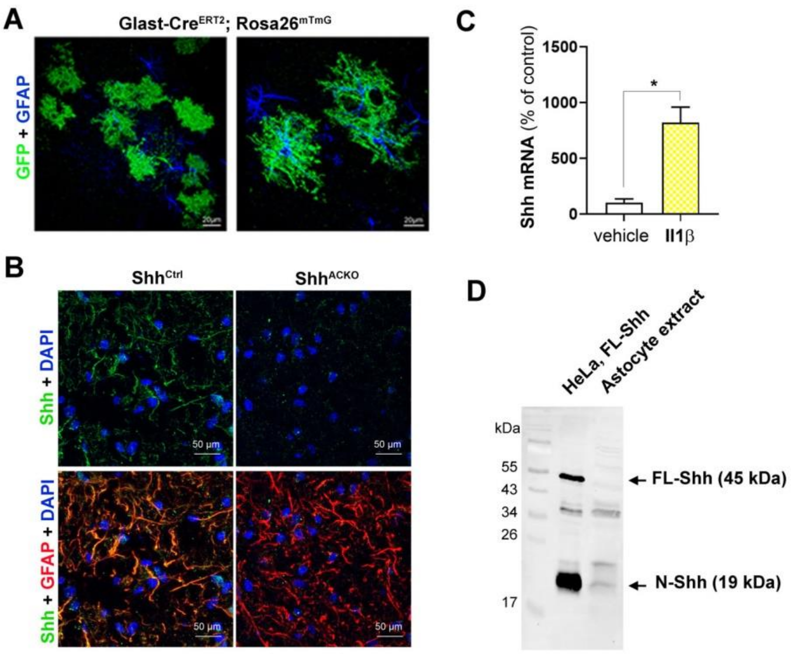


**Supplementary Figure 7:** (**A**) Spinal cord sections from GLAST-Cre$^{ERT2}$; Rosa26$^{mT/mG}$ were co-immunostained with anti-GFP (in green) and anti-GFAP (in blue) antibodies. The co-localization of GFAP and GFP staining validate the specific activity of the Cre recombinase within the astrocytes in the GLAST-Cre$^{ERT2}$ mouse model. (**B**) EAE was induced in both Glast-Cre$^{ERT2}$; Shh$^{Flox/Flox}$ (Shh$^{ACKO}$) and Shh$^{Flox/Flox}$ (Shh$^{ctrl}$) mice. Mice were sacrificed 32 days later. Spinal cord sections were co-immonstained with anti-Shh (in green) and anti-GFAP (in red) antibodies. (**C**) Cultured normal human astrocytes were treated with or not with 10 ng/mL Il1β for 24 hours. Shh mRNA expression was quantified by RT-qPCR. (**D**) Shh protein expression was analyzed by western blot in cell extract from both HeLa transfected with FL-Shh encoding plasmids and Il1β-treated astrocytes. *: $p \leq 0.05$. Mann Whitney test.
